# GRAMD2^+^ alveolar type I cell plasticity facilitates cell state transitions in organoid culture

**DOI:** 10.1101/2023.10.17.560801

**Authors:** Hua Shen, Weimou Chen, Yixin Liu, Alessandra Castaldi, Jonathan Castillo, Masafumi Horie, Per Flodby, Shivah Sundar, Changgong Li, Yanbin Ji, Parviz Minoo, Crystal N Marconett, Beiyun Zhou, Zea Borok

## Abstract

Alveolar epithelial regeneration is critical for normal lung function and becomes dysregulated in disease. While alveolar type 2 (AT2) and club cells are known distal lung epithelial progenitors, determining if alveolar epithelial type 1 (AT1) cells also contribute to alveolar regeneration has been hampered by lack of highly specific mouse models labeling AT1 cells. To address this, the *Gramd2^CreERT2^* transgenic strain was generated and crossed to *Rosa^mTmG^* mice. Extensive cellular characterization, including distal lung immunofluorescence and cytospin staining, confirmed that GRAMD2^+^ AT1 cells are highly enriched for green fluorescent protein (GFP). Interestingly, *Gramd2^CreERT2^* GFP^+^ cells were able to form organoids in organoid co-culture with Mlg fibroblasts. Temporal scRNAseq revealed that *Gramd2*^+^ AT1 cells transition through numerous intermediate lung epithelial cell states including basal, secretory and AT2 cell in organoids while acquiring proliferative capacity. Our results indicate that *Gramd2*^+^ AT1 cells are highly plastic suggesting they may contribute to alveolar regeneration.

## INTRODUCTION

The mammalian lung is comprised of more than 40 different cell types, of which a substantial portion includes epithelial cells (1). The cellular composition of the epithelial lining changes along the proximal-to-distal axis of the lung to meet different functional needs, ranging from mucociliary clearance to gas exchange (2). Many lung diseases including cancer and fibrosis, involve bronchiolar and alveolar epithelial cell dysfunction making it important to understand the mechanisms that regulate differentiation, maintenance, and repair of the lung epithelium. The most distal compartment of the lung, the alveolar epithelium, is comprised of alveolar epithelial type 1 (AT1) and type 2 (AT2) cells. AT1 cells are thin, flattened cells with attenuated cytoplasmic extensions that cover more than 90% of the alveolar surface and are responsible for gas exchange, whereas AT2 cells are cuboidal surfactant-producing cells located at the alveolar corners (3). There is strong evidence to support that during steady-state tissue maintenance and in response to injury, AT2 cells serve as the primary progenitors to both self-renew and differentiate into AT1 cells to regenerate the epithelial barrier (4–7). In contrast, AT1 cells have been viewed as terminally differentiated (8). However, more recently, the question of whether AT1 cells also possess a degree of plasticity has also been entertained.

In early studies, using a model in which rat AT2 cells in 2-dimensional (2D) culture lose their phenotypic hallmarks and acquire features of AT1 cells (so-called AT1-like cells), in one of the first demonstrations of AT1 cell plasticity, we showed that treatment with keratinocyte growth factor induced AT1-like cells to re-acquire AT2 cell characteristics, including lamellar bodies and expression of surfactant protein B, with reciprocal changes in AT1 cell markers (9). Similarly, we demonstrated that rat serum and alterations in cell shape (by switching cells cultured on attached collagen gels to detached collagen gels) partially reverse AT2 to AT1-like cell differentiation (10). In subsequent studies by others, rat AT1 cells were shown to proliferate and express both AT2 and club cell markers (SFTPC and SCGB1A1) in 2D culture (11, 12) and mouse HOPX^+^ PDPN^+^ AT1 cells formed organoids comprised of both AT1 and AT2 cells in 3 dimensional (3D) culture conditions (12). More recently, in lineage tracing studies, HOPX^+^ AT1 cells displayed the ability to give rise to SFTPC-expressing AT2 cells under specific injury conditions such as pneumonectomy and hyperoxic injury in adult mouse lungs (13). Additionally, it has been reported that mature AT1 cells can exit their terminally differentiated non-proliferative state and retract their cellular extensions by overexpression of SOX2 in *Hopx^CreER^* mice (14, 15). In contrast, lineage tracing studies of IGFBP2^+^ AT1 cells demonstrated the inability of this subpopulation of AT1 cells to differentiate into AT2 cells following pneumonectomy, suggesting that there may be a spectrum of plasticity across subpopulations of AT1 cells that may manifest differently depending on the specific Cre driver used for these studies (16). Despite demonstrations of AT1 cell plasticity *in vitro* and *in vivo*, in contrast to the large amount of evidence supporting the differentiation capacity of AT2 cells in repair following injury, studies of AT1 cell plasticity remain more limited. In large part this has been attributed to Cre driver lines generated using ‘classical’ AT1 markers such as AQP5, PDPN, HOPX, and AGER lacking specificity for AT1 cells in distal lung epithelium.

GRAMD2 is a GRAM-domain containing protein that is located at endoplasmic reticulum (ER)-plasma membrane (PM) junctions (17, 18). We recently identified GRAMD2 as a novel AT1 cell marker by comparing genome-wide expression profiles of purified rat AT2 and AT1 cells with multiple rat organs together with human and rat *in vitro* differentiated AT1-like cells (19). Immunofluorescence staining confirmed that GRAMD2 is expressed in AT1 cells and not present on the surface of AT2 cells in mouse lung (19). We recently developed *Gramd2^CreERT2^;mTmG* mice by crossing *Gramd2^CreERT2^*with *mTmG* reporter mice. Our initial characterization demonstrated green fluorescent protein (GFP) expression predominantly in AT1 cells, suggesting that these mice may be valuable for further exploring AT1 cell biology in health and disease (20). In this study, we extensively characterized the specificity of tamoxifen-mediated *Gramd2*-driven GFP expression in lungs of *Gramd2^CreERT2;mTmG^* mice and evaluated phenotypic plasticity of GFP^+^ AT1 cells during 3-dimensional (3-D) culture using immunostaining and single cell RNAseq (scRNAseq). We found that in *Gramd2^CreERT2;mTmG^* mice, GFP effectively labels AT1 cells. GFP^+^ cells were able to form organoids in 3-D co-culture and, surprisingly, AT1 cells can transit through multiple epithelial cell states, demonstrating that the GRAMD2*^+^* AT1 cell population has intrinsic plasticity and suggesting the capacity for regenerative renewal under specific environmental conditions.

## RESULTS

### *Gramd2^creERT2;mTmG^* mouse lineage labels AT1 cells in lung

To determine the specificity of tamoxifen-induced CreERT2 activation in our newly generated *Gramd2-creERT2* transgenic mouse strain, we performed lineage tracing by crossing the previously established reporter strain *Rosa^mTmG^* (21) with *Gramd2^creERT2^* mice, creating *Gramd2^creERT2;mTmG^*mice. Following tamoxifen induction, the specificity of GFP expression in *Gramd2^creERT2;mTmG^* mice was evaluated using protein lysates from several organs, including the lung (**Figure 1A**, **Figure S1**). GFP protein was detected specifically in the lung following tamoxifen treatment in both male and female mice, indicating lung-specificity of Cre-mediated recombination in *Gramd2^CreERT2;mTmG^*mice.

**Figure 1.**
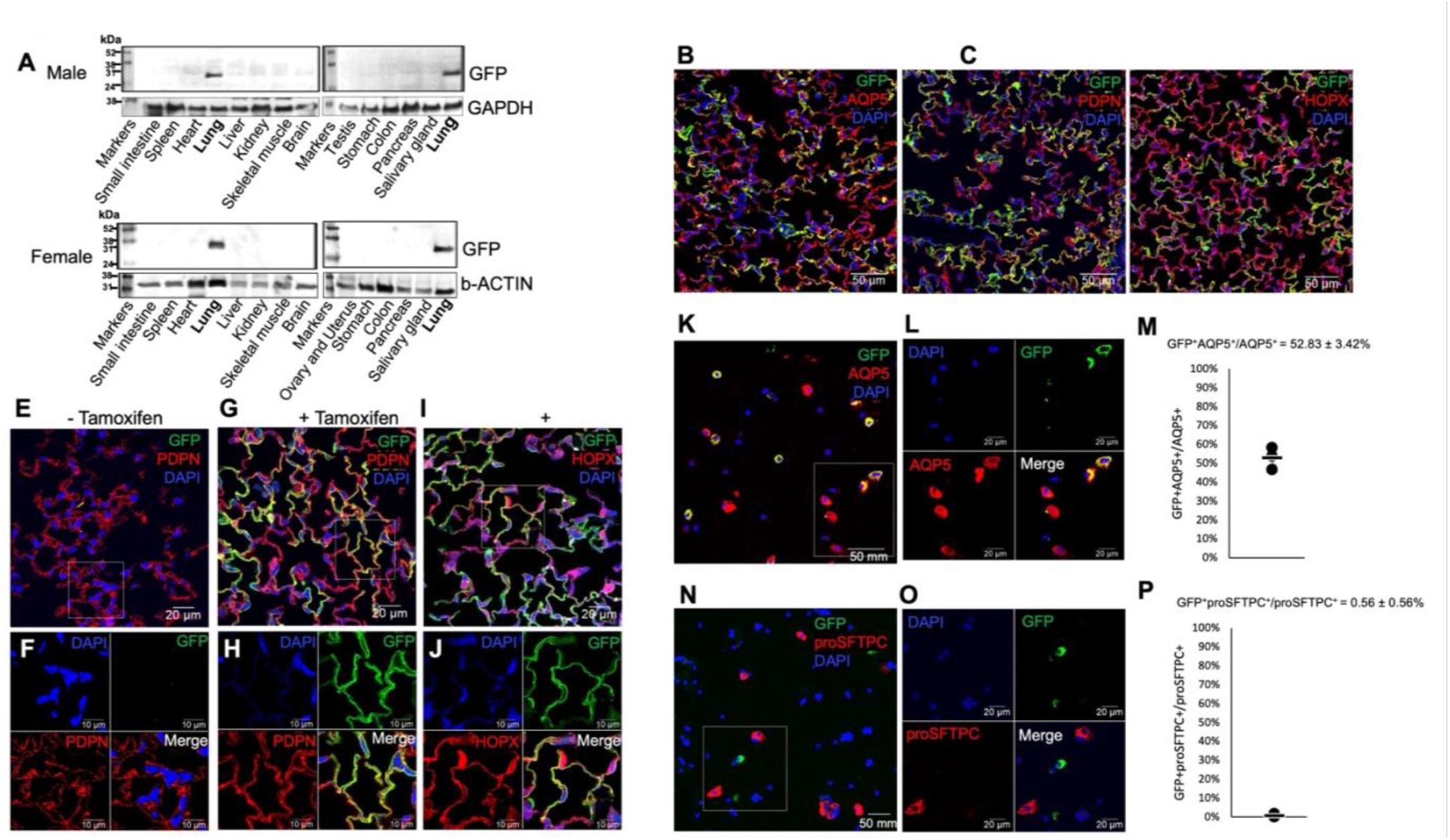
*Gramd2^creERT2;mTmG^* mouse lineage labels AT1 cells in lung. **A.** Representative western blots of GFP expression in different organs of male (top) and female (bottom) *Gramd2^CreERT2;mTmG^*mice. N = 4. **B.** Confocal tile scan of immunofluorescence staining in lung sections from *Gramd2^creERT2;mTmG^* mice. Green = GFP, red = AQP5, blue = DAPI. N=3. **C.** Confocal tile scan of immunofluorescence staining in lung sections from *Gramd2^CreERT2;^mTmG* mice. Green = GFP, red = PDPN, blue = DAPI. N=3. **D.** Confocal tile scan of immunofluorescence staining in lung sections from *Gramd2^creERT2;mTmG^* mice. Green = GFP, red = HOPX, blue = DAPI. N=3. **E.** Confocal imaging of lung sections from vehicle control (-tamoxifen) treatment of *Gramd2^creERT2;mTmG^* mice. Green = GFP, red = PDPN, blue = DAPI. N=3. Scale bar = 20 μm. **F.** Magnification of white box in (E). Green = GFP, red = PDPN, blue = DAPI. N=3. Scale bar = 10 μm. **G.** Confocal imaging of lung sections from tamoxifen-treated *Gramd2^creERT2;mTmG^* mice. Green = GFP, red = PDPN, blue = DAPI. N=3. Scale bar = 20 μm. **H.** Magnification of white box in (G). Green = GFP, red = PDPN, blue = DAPI. N=3. Scale bar = 10 μm. **I.** Confocal imaging of lung sections from tamoxifen treated *Gramd2^creERT2;mTmG^* mice. Green = GFP, red = HOPX, blue = DAPI. N=3. Scale bar = 20 μm. **J.** Magnification of white box in (I). Green = GFP, red = HOPX, blue = DAPI. N=3. Scale bar = 10 μm. **K.** Representative immunofluorescence staining of cytospins of crude cell preparations from distal lung tissue of *Gramd2^creERT2;mTmG^* mice. Green = GFP, red = AQP5, blue = DAPI. N=3. Scale bar = 50 μm. **L.** Magnification of white box in (K). Green = GFP, red = HOPX, blue = DAPI. Scale bar = 20 μm. **M.** Quantification of percentage of dual GFP^+^ AQP5^+^ cells present in crude cytospin preps from (K). Data represent the mean ± SEM of > 200 cells. Dots indicate percentage of cells for each sample and bar indicates the mean. N=3 independent preparations. **N.** Representative cytospin staining of crude cell preparations from distal lung tissue of *Gramd2^creERT2;mTmG^* mice. Green = GFP, red = proSFTPC, blue = DAPI. N=3. Scale bar = 50 μm. **O.** Magnification of white box in (N). Green = GFP, red = proSFTPC, blue = DAPI. Scale bar = 20 μm. **P.** Quantification of percentage of dual positive GFP^+^proSFTPC^+^ cells present in crude cytospin preparations from (N). Data represent the mean ± SEM of > 200 cells. Dots indicate percentage of cells for each sample and bar indicates the mean. N=3 preps.

Next, the specific populations of cells labeled with GFP by tamoxifen treatment of *Gramd2^creERT2;mTmG^* mice were determined by immunofluorescence co-localization of GFP with known cell type-specific markers. Negative staining in antibody isotype controls indicated that signal for known cell markers was specific (**Figure S2**). GFP signal in tamoxifen-induced *Gramd2^creERT2;mTmG^*lung sections extensively overlapped with several known AT1 cell markers, including AQP5, PDPN and HOPX. Interestingly, not all AQP5^+^, PDPN^+^ or HOPX^+^ AT1 cells in *Gramd2^creERT2;mTmG^* mice showed colocalization with GFP (**Figure 1B-1D**) (20). Higher magnification images further demonstrated colocalization of membranous GFP staining with PDPN^+^ and HOPX^+^ AT1 cells (**Figure 1E-1H**,) specifically under tamoxifen-induced CreERT2 activation conditions compared to no tamoxifen treatment (**Figure 1I, 1J**).

The above results suggest that the *Gramd2^CreERT2;mTmG^* mouse model can activate GFP in certain subpopulations of AT1 cells. To quantify the efficiency of CreERT2 recombination in AT1 cells and the relationship between *Gramd2^CreERT2;mTmG^*cell labeling and known AT1 cell markers, we isolated crude single cell preparations of distal lung tissue from *Gramd2^CreERT2;mTmG^* mice treated with tamoxifen and performed GFP and AQP5 dual immunofluorescence. The results showed that 52.83 ± 3.42% of AQP5^+^ cells were labeled with GFP (**Figure 1K-1M, Figure S3**). Consistent with AT1 cell specificity, GFP was expressed in only 0.56 ± 0.56 % of proSFTPC^+^ AT2 cells (**Figure 1N-1P**,). Additionally, a few GFP^+^ cells (some co-expressing SCGB1A1 and some acetylated tubulin (Ac-TUB)) were found in the airways (**Figure S4**). Collectively, these data indicate that GFP^+^ is enriched in AT1 cells in *Gramd2^CreERT2;mTmG^*mice.

### High yield AQP5^+^ AT1 cells can be isolated from *Gramd2^CreERT2;mTmG^* mice

Given the selective labeling of AT1 cells in distal lung epithelium of *Gramd2^CreERT2;mTmG^*mice, we explored whether this strain could be used to isolate AT1 cells for in vitro studies. Following our previously described protocol (22), the crude cell preparation yielded 6.74 X10^7^ ± 269,444 cells per mouse (N=5). FACS was used to perform negative selection using antibodies against CD31, CD34, and CD45 to remove endothelial, hematopoietic, and immune cells, respectively, and positive selection of GFP^+^ cells using a GFP Ab (**Figure S5A**). Cells from 129S6/SvEvTac mice were used as a GFP negative gating control (**Figure S5B**). To obtain a higher purity of AT1 cells, we adjusted the gate to ensure that only GFP^high^ cells were collected, and cells closely adjacent to the GFP^-^ population were excluded (**Figure S5**). Using this approach, we obtained 106,300 ± 11,077 GFP^high^CD31^-^CD34^-^CD45^-^ cells per mouse with > 90% viability in five independent preparations (total eight mice). The sorted GFP^+^ population was then stained with Abs against GFP, multiple AT1 cell markers (AQP5, PDPN, HOPX and IGFBP2) and AT2 cell marker proSFTPC. Compared to absence of detectable signal in negative controls (**Figure 2E**), IF and quantitation showed that among GFP^+^ cells, 98.43 ± 0.74% were AQP5^+^, 79.42 ± 5.54% were PDPN^+^, 75.61 ± 1.52% were HOPX^+^, and 54.51 ± 3.86% were IGFBP2^+^ (**Figure 2A-D, H**). Importantly, almost no overlap in signal was detected between GFP and proSFTPC (0.45 ± 0.26%), further confirming that CreERT2 is not activated in AT2 cells of *Gramd2^CreERT2;mTmG^* mice (**Figure 2F-H**). However, ∼ 1% GFP^+^Ac-TUB^+^ cells were detected (**Figure S6**). Together, these results indicate that GFP^+^ cells isolated by flow cytometry from *Gramd2^CreERT2;mTmG^* mice are highly enriched for subpopulations of AT1 cells.

**Figure 2.**
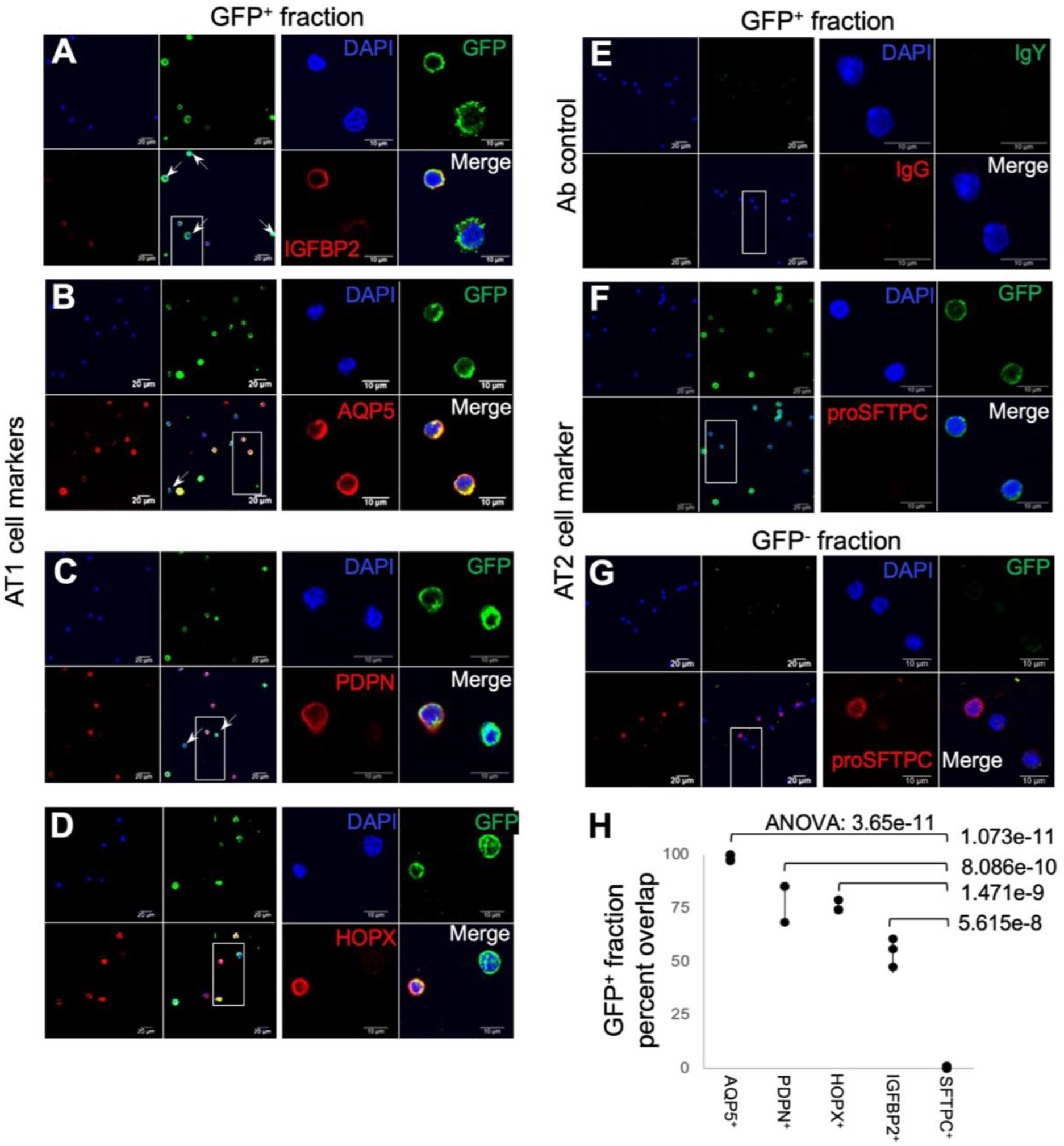
High yield AQP5^+^ AT1 cells can be isolated from *Gramd2^CreERT2;mTmG^* mice. **A-F.** Confocal images of cytospin staining of GFP^+^ FACS-sorted fraction of crude distal lung cell isolates. **A.** Green = GFP, red = IGFBP2, blue = DAPI. N=3. Scale bar (left) = 20 μm, (right) = 10 μm. **B.** Green = GFP, red = AQP5, blue = DAPI. N=4. Scale bar (left) = 20 μm, (right) = 10 μm. **C.** Green = GFP, red = PDPN, blue = DAPI. N=3. Scale bar (left) = 20 μm, (right) = 10 μm. **D.** Green = GFP, red = HOPX, blue = DAPI. N=3. Scale bar (left) = 20 μm, (right) = 10 μm. **E.** Ab control = primary antibody control lacking Fc recombinant region targeting antigen. Green = IgY isotype control, red = IgG isotype control, blue = DAPI. N=4. Scale bar (left) = 20 μm, (right) = 10 μm. **F.** Green = GFP, red = proSFTPC, blue = DAPI. N=4. Scale bar (left) = 20 μm, (right) = 10 μm. **G.** Confocal images of cytospin staining of GFP(^-^) FACS sorted fraction of crude distal lung cell isolates. Staining as in (F). N=4. Scale bar (left) = 20 μm, (right) = 10 μm. **H.** Quantification of overlap between known AT1 and AT2 cell markers and GFP signal in GFP^+^ FACS-sorted fraction of crude distal lung cell isolates using cytospin staining. Data are displayed as percent of cells with overlapping expression. Dots represent biological replicates. Signifcance was calculated between AT1 cell marker/GFP^+^ overlap and AT2 cell marker (proSFTPC) / GFP^+^ overlap using a one-sided ANOVA and Bonferroni false positive correction.

### GFP^+^ cells from *Gramd2^creERT2;mTmG^* mice form SFTPC^+^ organoids in 3-D co-culture

To investigate the ability of GFP^+^ AT1 cells from *Gramd2^CreERT2;mTmG^*mice to form organoids, we co-cultured sorted GFP^+^ cells and Mlg fibroblasts in Matrigel on Transwell filter inserts. We first evaluated the colony forming efficiency (CFE) of GFP^+^ cells from *Gramd2^creERT2;mTmG^* mice in organoid culture. As shown in **Figure S7**, organoid numbers increase as seeding density increases, while CFE remains similar (∼0.6%) (**Table 1**) across a range of 1,875-15,000 GFP^+^ cells plated per well. 15,000 GFP^+^ cells plated with 100,000 Mlg fibroblasts were used for all subsequent experiments. Under these conditions, GFP^+^ cells formed organoids which increased in size over time in culture from Day 6 through Day 25 (**Figure 3A**). To confirm that each individual organoid arises from a single seeded cell, we co-cultured 10,000 GFP^+^ AT1 cells from *Gramd2^CreERT2;mTmG^*mice and 10,000 RTM^+^ AT2 cells from Sftpc*^creERT2;RTM^* mice with 100,000 Mlg fibroblasts in Matrigel and found that even at this high seeding density (20,000 epithelial cells), AT1 and AT2 cells formed distinct organoids of either green or red color, with no mixed organoids being detected (**Figure S8**).

**Figure 3.**
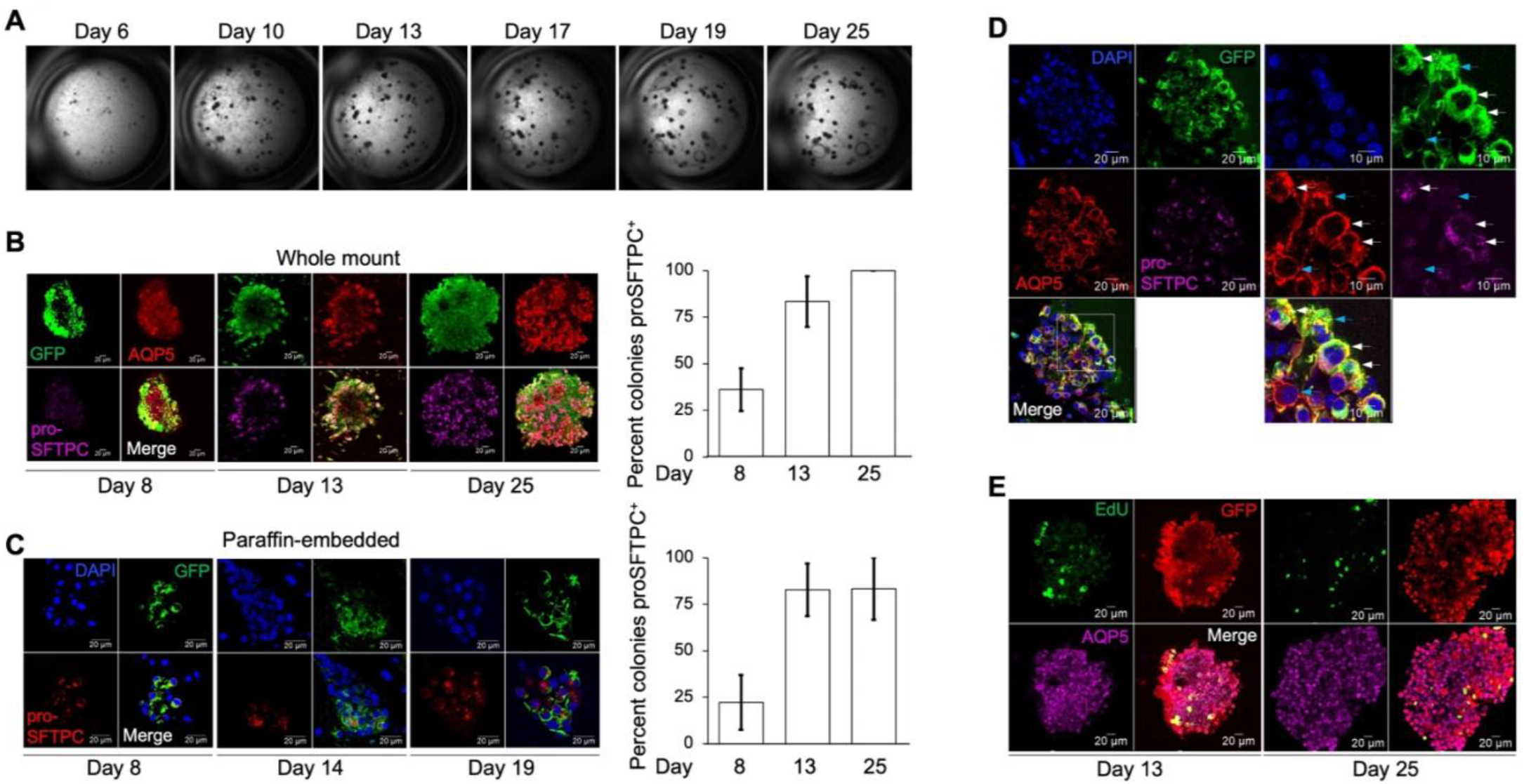
GFP^+^ cells from *Gramd2^creERT2;mTmG^* mice form SFTPC^+^ organoids in 3-D co-culture. **A.** Organoid formation of FACS-sorted GFP^+^ cells from *Gramd2^creERT2;mTmG^* mice co-cultured with Mlg fibroblasts in Matrigel. Day 6, Day 13, Day 17, Day 19 and Day 25 indicate the number of days in culture post-seeding in Matrigel. N=6. **B.** Representative whole mount confocal IF images (left) and quantification (right) of organoids formed in Matrigel from GFP^+^ *Gramd2^creERT2;mTmG^* cells. Green = GFP, red = AQP5, purple = proSFTPC. Scale bar = 20 μm. N=3. **C.** Representative confocal images (left) and quantification (right) of proSFTPC^+^GFP^+^ paraffin-embedded organoid sections. Green = GFP, red = proSFTPC, blue = DAPI. Scale bar = 20 μm. N=3. **D.** Representative confocal images of paraffin-embedded organoid sections co-stained for GFP (green), AQP5 (red), proSFTPC (purple), and DAPI (blue) on Day 19. Scale bar = 20 μm (left), 10 μm (right). White arrows indicate GFP^+^AQP5^+^proSFTPC^+^ cells, blue arrows indicate GFP^+^AQP5^+^proSFTPC^-^ cells. **E.** Representative whole mount confocal images of organoids immunostained for GFP (red), AQP5 (purple) and EdU (green) on Day 13 and Day 25 post-cell seeding. Scale bar = 20 μm. N=3.

To examine their differentiation potential, organoids formed from GFP^+^ cells were harvested on days 8, 13 or 14, 19, and 25 and analyzed by IF using either whole mount or paraffin-embedded sections. All organoids visualized using whole mount IF were GFP^+^, and the GFP^+^ cells in the organoids expressed AQP5 on days 8, 13, and 25 (**Figure 3B**). Isotype antibody controls were negative (**Figure S9**), indicating that staining for known cell markers was specific. Additionally, organoids visualized using whole mount IF were double positive for GFP and GPRC5A, another AT1 cell marker (23), on day 25 (**Figure S10**) indicating that some cells within each organoid still express AT1 cell markers at these later time-points in culture. Interestingly, despite a starting population of <1% proSFTPC^+^ cells, the percentage of proSFTPC^+^ cells detected in whole mount organoids increased by day 8. This trend continued at later time points, where an even greater percentage of proSFTPC^+^ cells was observed (**Figure 3B**, **Figures S11-13** (videos)). Quantification of organoids visualized by whole mount IF supported this observation, in that proSFTPC expression was seen in 36.11 ± 11.34 % of organoids on day 8, 83.33 ± 13.61% of all observed organoids on day 13, and 100 ± 0% of all organoids on day 25 (**Figure 3B**).

Similar findings were observed in paraffin-embedded organoid sections (**Figure 3C**), which allow more detailed visualization of cellular compartments (membrane and cytoplasm) than whole mount images. (Quantification of paraffin-embedded sections showed proSFTPC in 22.23 ± 14.7 % of organoids on day 8, 82.75 ± 14.22 % of organoids on day 14, and 83.33 ± 16.67 % of organoids on day 19 (**Figure 3C**). Triple staining of paraffin-embedded organoids with known AT1 and AT2 cell markers further showed that GFP^+^AQP5^+^SPC^+^ and GFP^+^AQP5^+^SPC^-^ cells existed within the same organoid (**Figure 3D**), indicating heterogeneity of cell identity within the organoids. To examine if the GFP^+^ cells proliferate, organoids were administered EdU 24 hours prior to harvest. On days 13 and 25, EdU signal was detected in some GFP^+^ AQP5^+^ cells by organoid whole mount IF (**Figure 3E**, **Figure S14**). This result suggests that GFP^+^ cells from *Gramd2 ^creERT2;mTmG^* mice have the ability to proliferate and differentiate into proSFTPC-expressing AT2 cells.

### Gramd2^+^ derived organoids contain specific subpopulations of AT1 cells

To further investigate if the GRAMD2^+^ AT1 cell population is distinct from other known AT1 cell populations, we triple stained GFP and proSFTPC with either HOPX or IGFBP2. HOPX^+^ cells have been reported to give rise to proSFTPC^+^ cells in 3-D culture and in vivo following pneumonectomy and hyperoxic injury (13, 15), while IGFBP2^+^ AT1 cells were unable to differentiate into AT2 cells in 3-D culture and were defined as terminally differentiated (16). We found that GFP^+^ (GRAMD2^+^) proSFTPC^+^ cells include both HOPX^+^ and HOPX^-^ cells (**Figure 4A**), suggesting that GRAMD2^+^ AT1 cells both overlap with and are distinct from HOPX^+^ cells. None of the GFP^+^ cells were IGFBP2^+^ (**Figure 4B**), suggesting that either IGFBP2^+^ AT1 cells do not give rise to organoids as previously reported (16) and/or that GRAMD2^+^ AT1 cells do not further mature into IGFBP2^+^ AT1 cells under these culture conditions.

**Figure 4.**
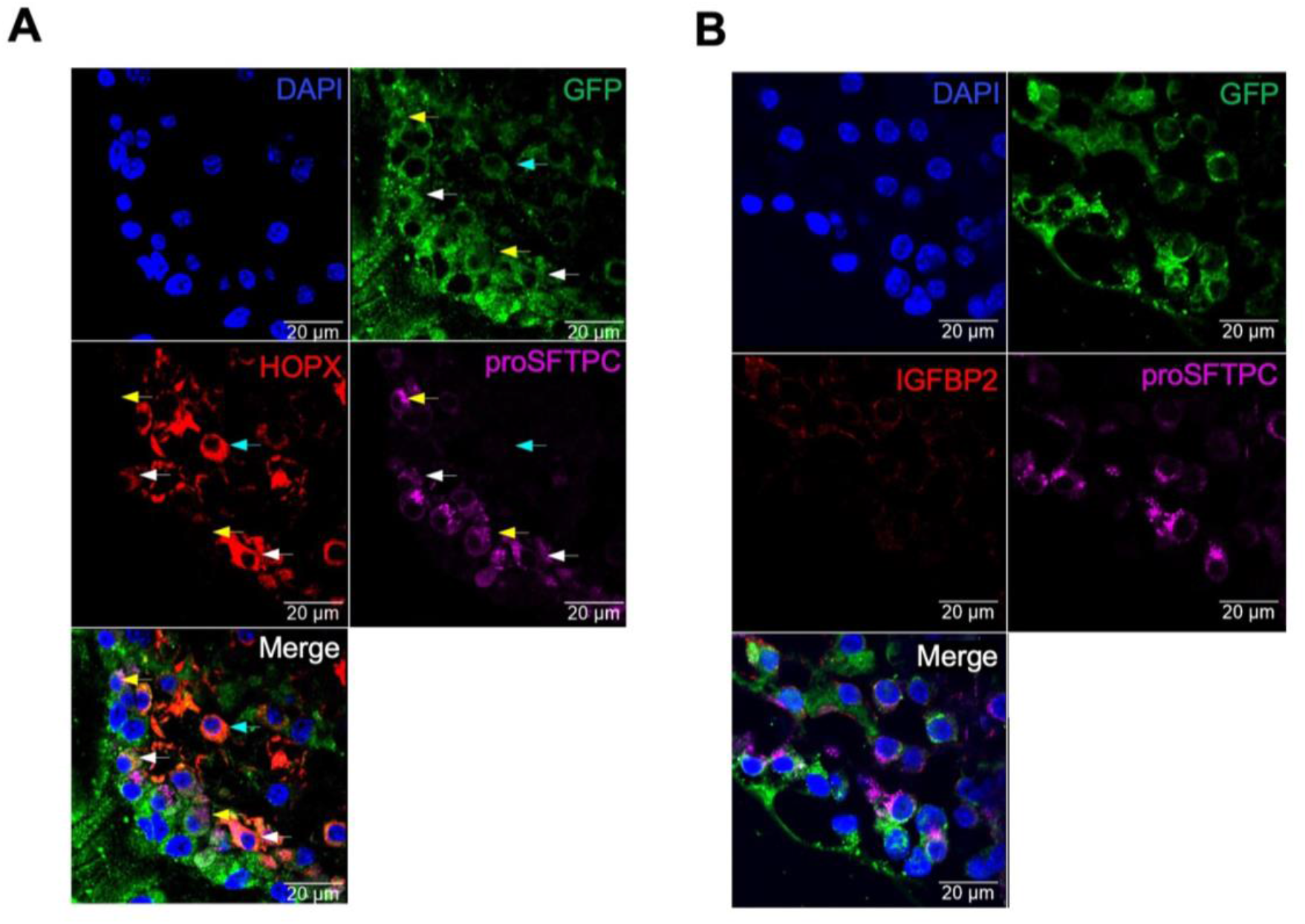
GRAMD2^+^ cell-derived organoids contain specific subpopulations of AT1 cells. **A.** Representative confocal IF image of organoids triple-stained for GFP (green), HOPX (red) and proSFTPC (purple) in paraffin-embedded section on Day 19. DAPI is the nuclear counterstain (blue). White arrows indicate GFP^+^HOPX^+^proSFTPC^+^ cells, yellow arrows indicate GFP^+^HOPX^-^ proSFTPC^+^ cells, and cyan arrow indicates GFP^+^HOPX^+^proSFTPC^-^ cells. Scale bar = 20 μm. N=3. **B.** Representative confocal IF image of organoid triple-stained for GFP (green), IGFBP2 (red), proSFTPC (purple) and DAPI (blue) in paraffin-embedded section on Day 19. Scale bar = 20 μm. N=3.

### Single cell RNAseq of GRAMD2^+^ cell-derived organoids reveals AT1 cell plasticity

To further characterize the differentiation trajectory of GRAMD2^+^ AT1 cells in 3-D culture, we undertook single cell RNAseq of sorted GFP^+^ cells on day 0 and organoids on days 8, 13 and 20 by scRNAseq (**Figure 5A**). Following quality control analysis to remove poor quality cells (**Figure S15**) and selection for EpCAM^+^ epithelial cells, we observed a distinct transition of cell transcriptomes over time in culture (**Figure 5B**). We also observed that EpCAM^+^ cells on D0 showed little to no proliferative capacity; however, by Day 8 and persisting through Day 13 in culture a distinct subset of cells had acquired proliferative capacity as determined by the presence of cells in S and G2/M phase of the cell cycle (**Figure 5C**). To understand the transcriptional changes coinciding with the observed increases in proliferation, we performed cell type identity analysis using known distal lung epithelial transcriptomic patterns present in the mouse LungMAP database. Strikingly, we observed that Day 0 cells consisted primarily of AT1 cells, and a smaller fraction of transcriptionally distinct ciliated cells (**Figure 5D**). However, during culture the cellular fates of these two populations varied dramatically. The ciliated population retained their ciliated identity and remained transcriptionally clustered with the Day 0 population throughout all measured timepoints. However, the AT1 annotated D0 population disappeared by Day 8. In its place, several distinct cellular identities were observed, including proliferative AT2 and secretory (club) cells, as well as cells expressing basal markers (KRT5 and KRT14) (**Figure 5D**). This pattern continued on Day 13, with distinct populations of proliferative AT2, secretory, and basal cells visible. Expression of KRT5 was confirmed in D14 organoids by immunostaining (**Figure S16**). By Day 20, the proliferative transcriptional signature dissipated, while cells annotated as differentiated AT2 cells increased dramatically in the overall percentage of cells observed (**Figure 5E**). Cells annotated as having AT1 cell transcriptional patterning also reappeared, although they failed to acquire markers of fully differentiated AT1 cells such as *Igfbp2*. To validate these findings, we reannotated the EpCAM^+^ population with both the human LungMAP and the Human Lung Cell Atlas version 2 (**Figure S17**), which demonstrated replication of basal, secretory, AT1, and AT2 cell identities, as well as the presence of a subset of ciliated cells on Day 0. Further interrogation of known cell type specific markers aligned with Azimuth-predicted cell identities (**Figure 5F**), consistent with GFP^+^ AT1 cells transitioning through multiple cellular states as organoids formed in 3-D culture.

**Figure 5.**
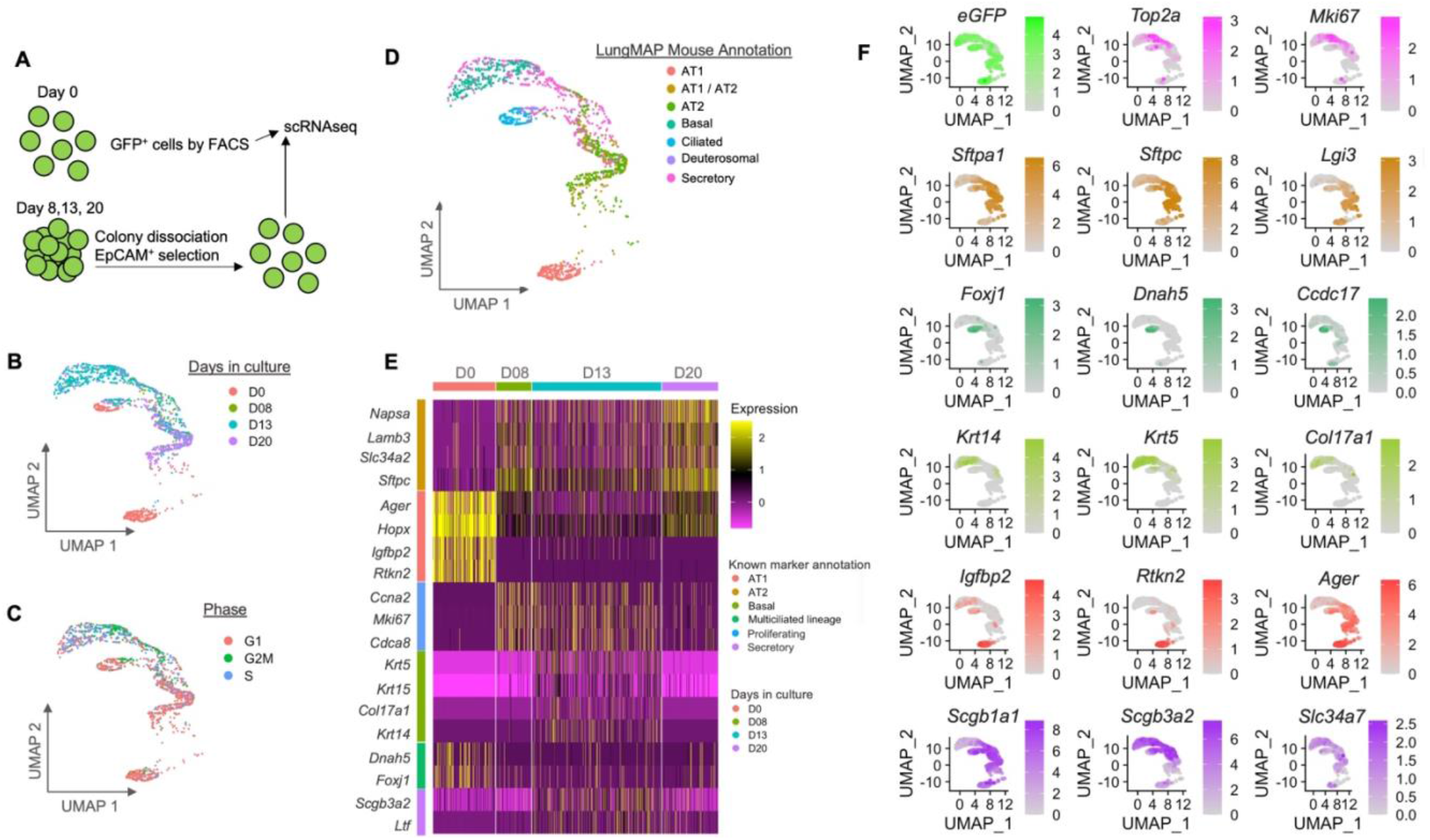
Single cell RNAseq of GRAMD2^+^ cell-derived organoids reveals plasticity of AT1 cells. **A.** Schematic showing single cells for scRNAseq obtained from *Gramd2^creERT2;mTmG^* mice using FACS and dissociation of cultured organoids using MACS. **B.** UMAP of EpCAM^+^ epithelial cell population, colored by days in culture; Day 0 (D0) = red, Day 8 (D08) = green, Day 13 (D13) = teal, and Day 20 (D20) = purple. **C.** UMAP of EpCAM^+^ epithelial cell population, colored by phase of the cell cycle: G1/G0 phase = red, S phase = pale blue, G2/M phase = green. **D.** UMAP of EpCAM^+^ epithelial cell population annotated to the mouse LungMAP reference using Azimuth. Alveolar epithelial type 1 (AT1) = red, alveolar epithelial type 2 (AT2) = gold, basal = peridot, multiciliated/ciliated cells = green, none/unassigned = teal, rare/neuroendocrine = blue, secretory/club cells = purple, submucosal secretory = pink. **E.** Heatmap of known markers for EpCAM^+^ epithelial cells present in the adult lung stratified by days in culture. Days colored as in (B). x axis = length of time FACS-sorted GFP^+^ AT1 cells were in culture; y axis = known cell markers for distinct lung cell populations. Cell markers colored as in (D). Heatmap is gradient colored by row. Z scale: purple = no expression, black = some expression, yellow = high expression. **F.** UMAPs of indicated cell type specific markers. UMAPs are colored as in (D) by their associated cell types.

### Pseudotime analysis of *Gramd2^CreERT2;mTmG^* organoids reveals intermediate cell states

Our observations of multiple cellular states arising from *Gramd2^CreERT2;mTmG^* GFP^+^ cells suggested that Gramd2^+^ AT1 cells are highly plastic and can transition through various lung states in organoid culture to facilitate proliferation and re-differentiation of distinct distal lung cellular populations. However, this did not explain how these transitions occurred or the relationship between observed cell states in 3D organoid culture. Therefore, we sought to understand how these cell fate transitions were occurring using trajectory analysis. CellRank was utilized to determine the temporal and transcriptomic relationships among the observed cell fate transitions (**Figure 6A**). Progression over time was then calculated using a real-time kernel, which relied on experimental timepoints of cells in culture (**Figure 6B**), as well as a pseudotime kernel, which relied on transcriptomic velocity (**Figure 6C**). Both methods showed cells of AT1 cell identity transitioning into the AT1/AT2 population and subsequently proceeding toward multiple lineages, including AT2, secretory, and basal (**Figure 6D**). To further deconvolute the relationship between cell fate transitions, we performed dimensional reduction to see root relationships between cell fates (**Figure 6E, Figure S18**). This revealed that after the AT1 population near-universally transitioned to the AT1/AT2 intermediate state, the differentiation hierarchy bifurcated into AT2 and secretory populations. Those cells that transitioned to the secretory fate largely continued to transition toward a basal identity. To understand what drove the observed cell fate transitions, we leveraged transcriptomic changes over time using both an unsupervised analysis, which extracted the top 10 genes with the largest fold changes in pseudotime while in organoid co-culture regardless of their known activities in lung biology to find key driver genes, and a supervised analysis, which utilized well-established markers of lung cell state transition. Unsupervised pseudotime analysis results were then plotted via heatmap to visualize their changes over time (**Figure 6G**). We observed that many of the genes predicted to be drivers of pseudotime in Gramd2^+^ At1 cells in organoid culture were genes with known association with alveolar cell type identity, including *Ager*, *Aqp4*, *Slc34a2*, *Napsa*, *Lgi3*, *Lamp3* and Sftpa2. We also identified several genes driving pseudotime changes that do not have well-described roles in alveolar biology but have been implicated in lung disease, including *Klk10*, *Klk13*, *Fxyd3*, and *Med24* (24–27). The previously described proliferation signal, including *Cdc8*, *Cenpf*, *Ube2c*, *Mki67*, and *Top2a* was also observed. Supervised analysis using known markers of AT1 to AT2 cell transition also showed a decrease in AT1 (**Figure 6H**) and increase in AT2 cell identity (**Figure 6I**).

**Figure 6.**
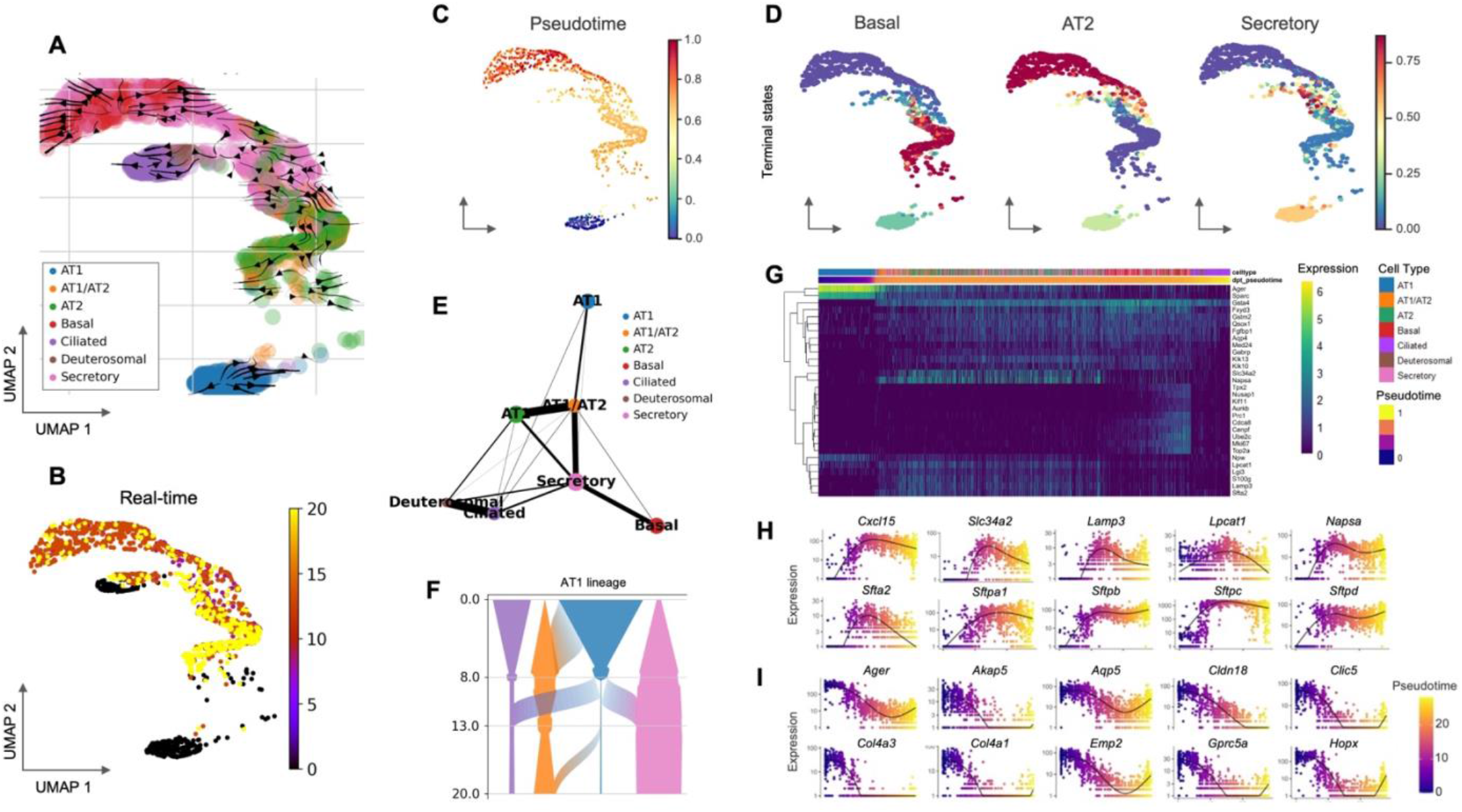
Pseudotime analysis of *Gramd2^CreERT2;mTmG^* organoids reveals intermediate cell states. **A.** UMAP of pseudotime velocity analysis. Arrows indicate direction of cell state transition. Colors indicate predicted cell types as annotated by Azimuth using the mouse LungMAP reference; blue = AT1, orange = AT1/AT2, green = AT2, red = basal, purple = ciliated, brown = deuterosomal, pink = secretory. **B.** UMAP of real-time trajectory analysis based on experimental timepoints. Black = Day 0, purple = Day 8, red = Day 13, and yellow = Day 20. **C.** Pseudotime assignment of cell states as initial/starting populations (blue) intermediate states (yellow/orange) and terminal states (red). **D.** Pseudotime assignment of terminal cell states as ending cell populations. High probability terminal states for the indicated cell populations = blue, low probability = red, probability is colored according to the color gradient between red to blue. **E.** Fate trajectory diagram of connection between transitional cell states. Colored circles indicate distinct transcriptomic patterns indicative of known cell identities present in the Mouse LungMAP reference database. Cell identities are colored as in (A). **F.** Fate transitions associated with cells of AT1 cell origin. Y-axis = days in culture, X-axis = proportion of cells in the indicated cell identity, grey swoops indicate proportion of cells transitioning from AT1 cell population to the indicated identities. Cell identities are colored as in (A). **G.** Heatmap of pseudotime-calculated gene intensity of the top 10 genes associated with cell fate transitions. Gene expression is scaled as a gradient from no expression (blue) to high expression (yellow). Cell identities are colored as in (A). Pseudotime is indicated by color gradient, initial timepoint (Day 0) = blue, to final timepoint (Day 20) = yellow. **H.** Pseudotime expression levels for individual genes associated with AT2 cell identity. Y-axis = expression level of indicated genes in scRNAseq cell population. Pseudotime is scaled as a color gradient; Day 0 (initial state) = blue, Day 20 (terminal states) = yellow. **I.** Pseudotime expression levels for individual genes associated with AT1 cell identity. Y-axis = expression level of indicated genes in scRNAseq cell population. Pseudotime is scaled as a color gradient; Day 0 (initial state) = blue, Day 20 (terminal states) = yellow.

The relationship between the ciliated cell population and the AT1 cell population was less clear. Ciliated cells on Day 0 largely transitioned to deuterosomal cell fate throughout the remaining timepoints (**Figure 6A**). Trajectory analysis revealed that there was a limited interconnection between primary AT1 cells and the deuterosomal cell state, likely due to the fact that deuterosomal cells arose from the intermediate basal cell state at later time points in organoid co-culture as they redifferentiated to airway cells fate (**Figure 6E**). Regardless, the ciliated population was not a major contributor to the observed AT1/AT2, AT2, secretory or basal cell populations (**Figure S19**).

### *Gramd2^CreERT2;mTmG^* associated transcriptional cell states are distinct from *Sftpc*- and *Trp63*-driven CreERT2 models

Our observations of multiple distinct cellular states arising from the GRAMD2^+^ AT1 cell population in distal mouse lung raised several questions regarding the uniqueness of these transitions to AT1 cells in organoid culture. To determine if similar cell fate transitions were observed with AT2 cells, we utilized previously published data on Sftpc*^CreERT2^* primary cells and 3-D organoid co-culture that had undergone scRNAseq on Day 14 (28) as well as primary basal cells isolated using the *Trp63^CreERT2^* transgenic model (29). We observed that both the primary AT2 cells derived from the Sftpc*^CreERT2^* model and primary basal cells derived from the *Trp63^CreERT2^* model showed related-but-distinct transcriptomic signatures from GFP^+^ cells from *Gramd2^CreERT2;mTmG^* cells (**Figure 7A**). The proliferative signature associated with AT2 proliferative cells was present in the Sftpc*^CreERT2^* but not in the *Trp63^CreERT2^*cell population (**Figure 7B**). Likewise, the proliferative basal population present in organoids derived from *Gramd2^CreERT2;mTmG^* GFP^+^ cells was associated with the subpopulation of proliferating basal cells present in *Trp63^CreERT2^* cells but not Sftpc^*CreERT2*^-derived cells (**Figure 7B**). Plotting each separate primary and organoid-derived sample from all three cells of origin allowed us to visualize how transcriptional states from all three distinct primary cells compared (**Figure 7C-D**). Annotation of the combined dataset using known markers of cell identity in mice revealed that AT2 cells also underwent increased proliferation in organoid culture, adoption of intermediate cell states, and differentiation to AT1-like cells (**Figure 7E**). This is consistent with what was observed previously (28). However, Sftpc*^CreERT2^* cells failed to acquire submucosal secretory and basal identities, highlighting how these two primary cell populations behave differently in organoid culture. This was further supported by comparing the relative abundance and levels of known marker gene expression across samples, with markers of basal cell identity arising at intermediate time points of *Gramd2*-but not *Sftpc*-derived organoids (**Figure 7F**). Taken together, our results indicate that GRAMD2^+^ AT1 cells can transition through multiple lung epithelial lineages in 3-D organoid co-culture in a manner distinct from SFTPC^+^ AT2 cells.

**Figure 7.**
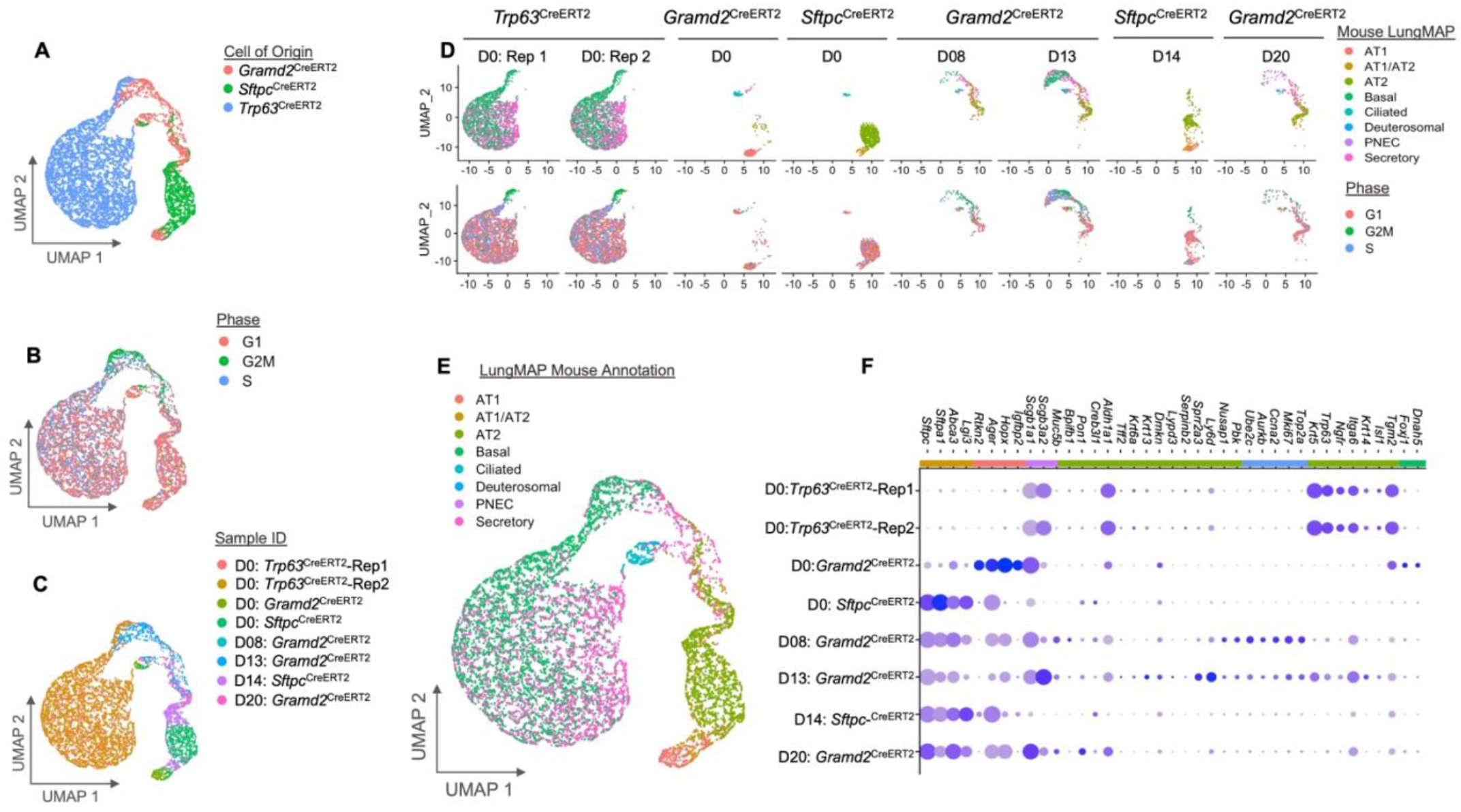
*Gramd2^CreERT2;mTmG^*associated transcriptional cell states are distinct from *Sftpc*- and *Trp63*-driven CreERT2 models. **A.** UMAP of transcriptomic profile relationships between EpCAM^+^ epithelial cells derived from *Gramd2^CreERT2^* (red), *S*ftpc*^CreERT2^* (green), and *Trp63^CreERT2^* (blue) reporter mice from either primary cell isolates or organoid cultures. **B.** UMAP of EpCAM^+^ epithelial cells derived from all three (3) CreERT2 models labeled by cell cycle phase. G1/G0 phase = red, S phase = blue, G2/M phase = green. **C.** UMAP of EpCAM^+^ epithelial cells colored by cell of origin and timepoint in organoid culture. *Trp63^CreERT2^* (basal) replicate 1 = red, replicate 2 = gold; *S*ftpc*^CreERT2^* (majority AT2 cells) D0 primary cells = green, D14 organoid culture = purple; *Gramd2^CreERT2^*(majority AT1 cells) D0 primary cells = peridot, D8 organoid culture = teal, D13 organoid culture = blue, D20 organoid culture = pink. **D.** UMAP of EpCAM^+^ epithelial cell population separated by cell of origin and timepoint in organoid culture. UMAPs are colored as in (C) by mouse LungMAP assigned cell identity in the top panel, and as in (B) by phase of the cell cycle below. Mouse LungMAP identity is colored as alveolar epithelial type 1 (AT1) = red, intermediate AT1/AT2 = orange, alveolar epithelial type 2 (AT2) = peridot, basal = green, ciliated = teal, deuterosomal = blue, pulmonary neuroendocrine cells (PNEC) = purple, secretory/club cells = pink. Phase of cell cycle is colored as in (B). **E.** UMAP of EpCAM^+^ epithelial cell population annotated to mouse LungMAP reference using Azimuth. **F.** Dot plot of known cell- and -phenotype specific markers for distal lung epithelial cell populations separated by cell of origin and time in organoid culture. Dot size is relative to percent of cells expressing the indicated cell marker; small dot = no cells express indicated gene, large dot = all cells express indicated gene. Color of dot represents gradient scale of expression level for the indicated gene within the given sample; no expression = grey, high expression = dark purple.

## DISCUSSION

AT1 cell function and plasticity have been far less studied than that of AT2 cells, largely due to the limited specificity of ‘classical’ AT1 cell markers for AT1 cell lineage tracing and fragility of AT1 cells which limits their isolation for in vitro studies. To address the need for additional specific AT1 cell markers, we identified a new lung-specific AT1 cell marker, GRAMD2 (19) through cross-species transcriptomic profiling across human, mouse and rat, and generated a new transgenic mouse line, *Gramd2^CreERT2^* (20). Here, we further characterized the AT1 cell specificity of GRAMD2 by crossing *Gramd2^CreERT2^* to *Rosa^mTmG^* reporter mice for lineage labeling of AT1 cells in distal lung epithelium and utilized these mice to investigate AT1 cell plasticity *in vitro*. In *Gramd2^CreERT2;mTmG^*reporter mice, by western blotting, GFP was detected specifically in lung. In distal lung, by IF, GFP was detected only in AT1, but not AT2 cells, with limited expression also in some airway epithelial cells, including ciliated and club cells. Intriguingly, sorted GRAMD2^+^ (GFP^+^) AT1 cells in 3-D culture formed organopids and proliferated, and differentiated towards AT2 and airway cells. Here, in the first scRNAseq analysis of AT1 cells in organoid culture, we show that GRAMD2^+^ AT1 cells have the capacity to transition through intermediate cell states, acquire proliferative capacity, and differentiate into several epithelial cell fates of lung identity. Taken together, these data indicate that the GRAMD2^+^ population of AT1 cells is highly plastic and suggest that *Gramd2^creERT2;mTmG^* mice may be used for enrichment of transiently differentiating AT1 cells (30, 31).

Several animal models have been generated to target AT1 cell-specific gene deletion and lineage tracing using Cre recombination in AT1 cells including *Aqp5^Cre^* (32), *Ager^Cre^* (33), *Hopx^Cre^* (34), *Rtkn2^Cre^* (35) and *Igfbp2^Cre^* (16). *Rtkn2^Cre^* and *Igfbp2^Cre^*models have been previously demonstrated to have a high degree of specificity for fully differentiated AT1 cells (16). In contrast, other widely used AT1 cell labeling models activate in a broader spectrum of cell types, depending on the mouse strain. Specifically, AQP5 expression is only entirely AT1 cell-specific in the 129S6/SvEvTac background, but consistently labels a small subpopulation of AT2 cells in the C57Bl6 background (32). *Hopx^Cre^* is expressed in AT2 cells and intermediate AT1/AT2 transitional cells (31, 36, 37). *Ager*^Cre^ was reported to label many AT2 cells (15). A recent paper also showed the expression of IGFBP2 in AT2 cells (38).

To date, only *Hopx*^Cre^ and *Igfbp2*^Cre^ have been used in studies to evaluate AT1 cell plasticity. Consistent with the notion that IGFBP2^+^ cells represent the most mature subpopulation of AT1 cells and are incapable of differentiation into AT2 cells, HOPX^+^/IGFBP2^+^ cells derived from the *Igfbp2*^Cre^ model lack the capacity to form organoids in 3-D culture (16), unlike HOPX^+^ / IGFBP2^-^cells which are able differentiate into AT2 cells (13, 15). Interestingly, initial populations of primary isolated GRAMD2^+^ AT1 cells contain both HOPX^+^ and HOPX^-^ cells, suggesting incomplete overlap between the GRAMD2^+^ and HOPX^+^ AT1 cell populations. Ciliated cells were also present in the primary isolated GRAMD2^+^GFP^+^ FACS isolated cell population, which was confirmed using staining for ciliated markers on tissue sections (**Figure S6**). A subpopulation of ciliated cells has been observed in multiple transgenic alveolar-cell labeling models, including our reanalysis of previously published TdTomato^+^ cells isolated from *S*ftpc*^CreERT2^*;Rosa*^TdTomato^* isolations (**Figure 7D**). This suggests that at least a subpopulation of ciliated cells also express multiple established alveolar epithelial cell markers.

In this study, we also found that GRAMD2^+^ AT1 cells were also able to proliferate and differentiate into multiple lung epithelial cell lineages in 3-D culture, including AT2, secretory, and basal identities. Strikingly, these results indicate that GRAMD2^+^ AT1 cells also exhibit plasticity and have the capacity to differentiate. Trajectory analysis revealed that two major fates arose from the GRAMD2^+^ AT1 cells placed in 3-D culture; those that transitioned through an AT1/AT2 intermediate cell state and terminated in an AT2 cell identity, and those that initially transitioned through an AT1/AT2 state and continued through a secretory/club cell state before differentiating further into a basal cell identity (**Figure 6E**). While HOPX^+^ AT1 cells were shown to have the ability to differentiate toward an AT2 identity, it remains to be seen if the HOPX^+^ AT1 cell population can also differentiate into more proximal lung epithelial identities, or if their transitional cell state capacity is limited to the AT1-to -AT2 branch we observed in a subset of the GRAMD2^+^ AT1 cell lineage (**Figures 6, S19**).

Interestingly, our reanalysis of Sftpc^+^ AT2 cells placed in 3-D organoid co-culture (28) revealed stark differences in the plasticity of AT1 and AT2 cells. While *Gramd2^CreERT2^*-derived AT1 cells shifted into intermediate cell states expressing proliferative, basal, and secretory cell markers, Sftpc*^CreERT2^*-derived cells either remained AT2 cells or differentiated further toward an AT1 cell identity; they did not differentiate toward a more proximal epithelial lung identity (**Figures 7D, 7F**). This could be explained by either differences in co-culture conditions, the fraction of AT2 cells represented in the *Sftpc^+^* population, or an inherent difference in the plasticity potential of AT1 vs AT2 cells in 3-D co-culture.

Multiple diseases of the distal lung display defects in differentiation and acquisition of aberrant transcriptional patterns. One example of this is idiopathic fibrosis (IPF), that presents with fibrotic encroachment throughout the parenchyma and eventual solidification of the lung. Cellular studies on IPF have revealed that extensive remodeling of the alveolar epithelium occurs, replacing normal alveolar populations with *Tp63^+^ Krt17^+^ Lam3^+^ Krt5^-^* aberrant basaloid cells (36), *Krt5*^+^ cells, dysregulated and expanded club and serous cell populations, with accompanying extensive “bronchiolization” (39). Multiple studies have postulated the starting population of these aberrant cells in diseased lungs, including AT2 cells and basal cells. Our results suggest that GRAMD2^+^ AT1 cells have the capacity to transition to a basaloid identity under specific conditions in 3-D organoid co-culture and may therefore also contribute to the dysregulated cellular phenotypes observed in IPF. GRAMD2^+^ AT1 cell plasticity in 3-D culture supports the possibility that GRAMD2^+^ AT1 cells may also contribute to distal lung regeneration following injury. In future studies, we plan to explore whether GRAMD2^+^ AT1 cells can regenerate the lung in models previously used to explore AT1 and AT2 cell plasticity such as hyperoxia and pneumonectomy (13, 15).

Another major disease that effects the distal lung is lung adenocarcinoma (LUAD). Activation of Kras, a known oncogenic driver, in HOPX^+^ cells can produce tumors with adenocarcinoma histology (13, 15). Our recent paper utilizing the *Gramd2^CreERT2^* model to activate Kras^G12D^ oncogenic driver expression in GRAMD2^+^ AT1 cells (20), showed that the resulting tumors had varied histology, including precancerous atypical adenomatous hyperplasia (AAH), lepidic, and papillary adenocarcinoma, which may reflect the AT1 transitional cell state responses to the stress induced by Kras^G12D^ activation. This is supported by observations in GRAMD2^+^ AT1 LUAD, where IF staining showed tumors were *SCGB3A2^+^ and KRT8*^+^ (20), markers which were observed in our 3-D organoid co-culture of GRAMD2^+^ AT1 cells undergoing cellular fate transitions. These data support that GRAMD2^+^ AT1 to AT2 cell differentiation could be impaired leading to accumulation of abnormal transitional cells in response to oncogene KRAS-meditated stress and suggests that induction of aberrant transitional cells during GRAMD2^+^ AT1 to AT2 cell differentiation in response to stressed stimuli could also be one of the mechanisms contributing to the pathogenesis of IPF.

Our choice of GRAMD2 for development of an AT1-labeling transgenic model for studying AT1 cell biology was based on our previous study identifying GRAMD2 as a highly enriched AT1 cell marker (19), independent of its cellular function. GRAMD2 is a GRAM domain-containing protein which is found in complex with other membrane-associated proteins (40). GRAMD2 localizes to the membrane contact sites (MCSs) between the endoplasmic reticulum and plasma membrane (ER-PM). In eukaryotic cells, the MCSs between the ER-PM are highly conserved and involved in lipid metabolism and Ca2^+^ signaling pathways (18). Interestingly, GRAMD2 is implicated in store-operated calcium entry (SOCE) mediated-metastatic processes (41, 42). In identifying GRAMD2 as an AT1 cell marker and developing the *Gramd2^CreERT2^* model to study GRAMD2^+^ AT1 cell biology, we may have inadvertently selected for a highly plastic subpopulation of AT1 cells with a functional role in disease progression. Further exploration of the functional role GRAMD2 plays in AT1 cell homeostasis and disease will further illuminate the behavior of distinct AT1 cell populations.

In summary, our study demonstrates that the GRAMD2^+^ AT1 population transitions into multiple distinct cellular fates in 3-D organoid co-culture. Identifying transitional cells with basal cell identity in GRAMD2-derived organoids may improve our understanding of the contribution of AT1 cells to lung repair after injury and pathogenesis of the disease (e.g., IPF). *Gramd2^CreERT2^* mice are therefore a valuable model to further elucidate the mechanisms underlying the formation of intermediate cell states and how these cells differentiate to repopulate distal lung cell populations in normal physiological conditions and following injury.

## MATERIALS AND METHODS

### Mice

*Gramd2*-*CreERT2;^mTmG^* mice were generated by breeding *Gramd2^creERT2^* with *mTmG* mice as previously described (20). Briefly, *Gramd2^creERT2^* mice were maintained in the 129S6 strain by backcrossing to 129S6/SvEvTac mice for 10 generations (Taconic Biosciences, Inc, San Diego, CA). For detection of the lineage label, *Gramd2 ^creERT2^* mice were crossed to *Rosa^mTmG^*mice (Jackson Laboratories, Bar Harbor, ME; Stock #007576) also on the 129S6/SvEvTac background. *Gramd2^creERT2^* and *mTmG* double heterozygous mice (*Gramd2^creERT2;mTmG^*) were used for evaluation of sites of GFP expression in various organs. *Gramd2^creERT2;mTmG^*mice were administered 200 mg/kg tamoxifen intraperitoneally on days 1, 3, and 5, and lungs were harvested seven days after the last tamoxifen injection. *S*ftpc*^creERT2;RTM^* mice were generated by crossing *S*ftp^creERT2^ mice (from H. A. Chapman, University of California, San Francisco) with *Rosa^mTmG^*reporter mice (21). All animal protocols were approved by the Institutional Animal Care and Use Committee (IACUC) at the University of Southern California.

### Western analysis

Dissected mouse tissues from the indicated organs were immediately transferred into protein lysis buffer as previously described (46). Briefly, mouse organs were homogenized in 2% SDS lysis buffer containing protease inhibitor mixture III (# 535140, Calbiochem, Billerica, MA) and phosphatase inhibitors (#P2850, Sigma) to obtain protein extracts. 40-80 μg protein was separated on SDS-PAGE gels and transferred to polyvinylidene difluoride (PVDF) membranes (#162–0177; Bio-Rad). Membranes were blocked in phosphate buffered saline (PBS)-Tween with 5% non-fat dry milk. Antibody (Ab) against GFP (#A6455; Thermo Fisher Scientific, Waltham, MA) was used at 1:1000 dilution. Bound primary Ab was visualized using secondary Ab conjugated to horseradish peroxidase (#sc-2030,1:2000, Santa Cruz Biotechnology, Inc, Dallas, TX) and chemiluminescent substrate (ECL Plus; # 32134; Thermo Fisher Scientific). Beta-actin (#sc-47778, 1:1000, Santa Cruz, Dallas, Texas) or GAPDH (#AM4300, 1:1000 Ambion, Waltham, MA) were used as loading controls.

### Preparation of crude cytospins

Lungs were digested by instilling 0.8 mg/ml collagenase/dispase (#10269638001 Roche, Indianapolis, IN) and 2 mg/ml pronase (#10165921001, Roche) via the trachea and incubating for 30 minutes at 37°C. Lung lobes were dissected away from the major airways, minced, and suspended in Dulbecco’s Modified Eagle Medium:Nutrient Mixture F-12 (DMEM/F12, #D6421, Sigma-Aldrich, St. Louis, MO) containing 0.01% DNase (# 10104159001, Roche). 100 µm and 40 µm filters (#352360 and #352340, Falcon, Glendale, Arizona) were used sequentially to remove debris. Erythrocytes were removed using red blood cell (RBC) lysis buffer (#11814389001, Roche). Cells were counted and trypan blue was used to measure viability. After counting, cells were fixed in 4% paraformaldehyde (PFA) for 10 minutes, washed with PBS, resuspended in PBS-FGH (2% fetal bovine serum (FBS), glucose 45% and HEPES 1M) buffer and the concentration adjusted to 1X10^5^ cells in 500 ml for each cytospin. Cytospins were prepared using EZ single cytofunnels (#1439045, Thermo Fisher Scientific) and a Shandon CytoSpin III cytocentrifuge (GMI, Ramsey, Minnesota). Cytospin slides were stored at -20°C until use.

### Fluorescence activated cell sorting (FACS)

Crude cell preparations from *Gramd2 ^creERT2;mTmG^* mouse lungs (isolated as above for cytospins) were incubated with primary Abs on ice for 30 minutes. CD31 (#13-0311-85, Thermo Fisher Scientific), CD34 (#14-0341-82, Thermo Fisher Scientific) and CD45 (#13-0451-85, Thermo Fisher Scientific) Abs were used to remove immune cells. GFP Ab (#A6455, Thermo Fisher Scientific) was used to select GFP^+^ cells. After washing with PBS-FGH three times, cells were incubated with secondary Abs for 30 minutes on ice. Ab-bound cells were sorted by FACS and data were collected using a BD FACSDiva8.0.2 system (BD Bioscience, La Jolla, CA). DAPI staining was used to exclude dead cells. The same crude cell processing method was used to collect cells from wild type 129S6/SvEvTac mouse lungs which were used as a negative gating control for GFP fluorescence for FACS.

### Organoid culture and preparation for immunofluorescence and scRNAseq

Sorted GFP^+^ cells (12,000–18,000) were seeded with 100,000 Mlg fibroblasts (#CCL 206, American Type Culture Collection, Manassas, VA) in growth factor-reduced Matrigel (#354230, Corning, Glendale, AZ) on Transwell filter inserts (#3470, Corning) in Dulbecco’s DMEM/F-12 (#D6421, Sigma-Aldrich) supplemented with 10% fetal bovine serum (FBS) (#SH30070.03, Hyclone, GE Healthcare Life Sciences, Marlborough, MA), 10 mg/l insulin, 5.5 mg/l transferrin, 2 mM glutamine, 100 units penicillin/streptomycin and 0.01 mM SB431542 (#1614/1, R&D systems). Organoids were harvested on days 6–25 of culture. Organoids grown in Matrigel on Transwell membranes were fixed in 4% PFA overnight at 4°C. The next day, membranes were removed and organoids and Matrigel pre-embedded in 100-150 µl Histogel (#22-110-678, Fisher Scientific), dehydrated in ethanol and then embedded in paraffin. Alternatively, organoids were used for scRNAseq as described below.

### Immunofluorescence (IF) staining

Lungs were perfused with PBS followed by inflation with 4% PFA at 25 cm H_2_O pressure, lobes were separated, and subsequently incubated in 4% PFA at 4°C overnight. Lungs were then dehydrated in ethanol and embedded in paraffin. For paraffin embedded lung sections and organoids from 3D culture, after de-paraffinization, sections (5 µm) underwent antigen retrieval using antigen unmasking solution (# H3301, Vector Laboratories, Burlingame, CA). Sections were washed and permeabilized with PBS containing 0.1% Triton X-100. Cytospin slides were first allowed to reach room temperature and then fixed with 4% PFA for 10 minutes. Sections or cytospin slides were blocked with CAS block (#008120, Thermo Fisher Scientific) at room temperature for 30 minutes to 1 hour, then incubated in primary Abs at 4°C overnight. Primary Abs were as follows: GFP (1:200-500, #13970, Abcam, Eugene, OR or #A6455, Thermo Fisher Scientific), AQP5 (1:200, #Aqp-005, Alomone Labs, Jerusalem 9104201, Israel), proSFTPC (1:200-500, #WRAB-SPC-9337, Seven Hills, Cincinnati, OH), PDPN (1:200 #127403 8.1.1, Biolegend, San Diego, CA), SCGB1A1 (1:500, #S9772, Santa Cruz), SCGB3A2 (1:200, #mUGRP1 AF3, 465, R&D, Santa Fe Springs, CA), SOX2 (1:100, #4900, Cell Signaling, Danvers, MA), HOPX (1:50, #SC398703, Santa Cruz), acetylated anti-tubulin (Ac-TUB, 1:200, #T6793, Millipore), IGFBP2 (1:50, #SC25285, Santa Cruz) and GPRC5A (1:200, #abx005719, Abbexa, Sugar Land, TX). Slides were mounted with media containing 4′,6-diamidino-2-phenylindole (DAPI) or propidium idodide (PI). Fluorescent (Leica DMI8, Leica Microsystems, Buffalo Grove, IL) and confocal (1 μm optical sections, Leica TCS SP8 confocal) images were captured at the same settings and at the same time for all sections within an experimental group.

### Whole mount immunofluorescence staining and EdU labeling of organoids

Whole mount staining of organoids was performed as previously described (47). Briefly, on days 8 (Day 8), 13 (Day 13), 19 (Day 19) and 25 (Day 25) of culture, membranes from Transwell inserts were transferred to a 10 cm petri dish. Organoids were gently isolated from Matrigel under a dissecting microscope and transferred to 1.5 ml tubes. TrypLE express (#12604-013, Gibco, Waltham, Massachusetts) was added to the tube and samples incubated at 37°C for 5 minutes to remove residual Matrigel. Organoids were then washed with PBS and transferred to 48-well culture plates. After fixation in 4% PFA for 45 minutes at 4°C, organoids were incubated in primary Abs at 4°C overnight. Primary Abs were as follows: GFP (1:200, #A6455, Thermo Fisher Scientific), AQP5 (1:50, #SC-9890, Santa Cruz), proSFTPC (1:200-500, #WRAB-SPC-9337, Seven Hills), SCGB1A1 (1:500, #S9772, Santa Cruz), acetylated tubulin (1:200, #T6739, Millipore) and KRT5 (1:200, #905903, Biolegend). For Edu labelling, organoids were treated with 10 μM 5-ethynyl-2’-deoxyuridine EdU (#E10187, Invitrogen) 24 hours prior to harvest. Click-iT® Plus EdU Alexa Fluor^TM^ 488 imaging kit (#C10637, Thermo Fisher Scientific) was used for EdU examination. After adding secondary Abs, organoids were placed on slides in clearing solution for imaging. Z-stack images (5 μm increments) were captured using a confocal microscope (Leica TCS SP8) and exported as TIFF images or videos.

### Single cell RNA sequencing (scRNAseq)

Primary isolates of FACS sorted GFP^+^ cells were defined as Day 0. On days 8 (Day 8), 13 (Day 13), and 20 (Day 20) of 3D culture, to obtain single cells, organoids were dissociated with dispase (#354235, BD Gibco) and TrypLE express (#12604013, Gibco). Biotinylated EpCAM Ab (#130-096-419, Miltenyi Biotec, San Diego, California), anti-biotin microbeads (#130-900-485, Miltenyi Biotec,) and MS columns (#130-042-201, Miltenyi Biotec) were used to enrich for epithelial cells. For each time point, cells were counted, and viability was measured using an automated cell counter (Countess 3, Thermo Fisher Scientific). Single cell suspensions were processed using Chromium Single Cell 3’ Reagent Kits v3.1 (10x Genomics, PN-1000128, PN-1000127 and PN-1000213). Briefly, cells resuspended in the reaction mix were loaded into the Chromium Single Cell Chip G, together with the 10x Barcoded Gel Beads and partitioning oil. The Chip G was placed into the 10x Chromium controller for cell partitioning, targeting a final number of 10,000 cells per sample. After partitioning, the GEMs (Gel Bead-In EMulsions) were transferred to the C1000 Touch Thermal Cycler for the first phase of reverse transcription followed by library preparation following the manufacturer’s instructions. Libraries were deep sequenced using the Illumina NovaSeq platform.

### Single cell RNAseq analysis

Raw FASTQ sequencing files were aligned to the mm10 genome that had been compiled with the addition of *mTmG* transgenes (21) using the Cell Ranger pipeline (Count v7.0.1). Aligned and filtered H5 files from all cells of origin were imported into R studio (v 2023.03.0+386) and analyzed using the Seurat package (v 4.3.0) built under R (v 4.3.0). Poor quality cells were filtered out by removing any cell with >5% mitochondrial contamination. The dataset was further filtered to remove non-epithelial cells by requiring >1 count of *Epcam* expression. Data were normalized, scaled, and clustered using the Seurat pipeline under a resolution value of 0.05. Annotation of known cell identities was performed using the LungMAP Consortium mouse reference using the online Azimuth interface. Secondary annotation of cell identities was performed using scRNAseq data generated by the Human Lung Cell Atlas version 2 (HLCAv2) present in Azimuth (48), which was built from several previously published scRNAseq studies (37, 49–55). Cells annotated as a cell type other than epithelial, were removed from further analysis. Heatmaps (Do.Heatmap), Dot plots (DotPlot), and UMAPs (FeaturePlot, DimPlot) were generated using Seurat in R with default color settings.

### scRNAseq trajectory analysis

To estimate differentiation direction of the cells placed in 3D organoid co-culture across multiple time points, the python package CellRank was used (56). First, cell-cell transition probabilities were estimated to create a Markov transition matrix. Using the CellRank RealTimeKernel, cells were matched within- and across-experimental time points by using the mathematical approach of optimal transport (57). To correct for cell growth and death within- and across experimental time points, proliferation and apoptosis markers were used. Second, the cell-cell transition matrix was used to identify initial, intermediate, and terminal macrostates and to compute fate probabilities using the Generalized Perron Cluster Cluster Analysis (GPCCA) algorithm (45).

### Data accessibility

Raw and processed data files for primary GFP^+^ AT1 cells (Day 0) and organoid-derived cells (Day 8, Day 13 and Day 20) from *Gramd2^CreERT2;mTmG^* mice are publicly available through the National Center for Biotechnology Information (NCBI) Gene Expression Omnibus (GEO) under identifier GSE235217. Data for AT2 primary (Day 0) and organoid-derived cells (Day 14) from *S*ftpc*^CreERT2;TdTomato^*mice were generated previously (28). Data from primary basal cells from *Trp63^CreERT2^* reporter mice were previously published (29). Initial annotation of organoid cell identity was performed using the LungMAP Consortium (U01HL122642) using the Azimuth interface for human (https://app.lungmap.net/app/azimuth-human-lung-cellref-seed) and mouse (https://app.lungmap.net/app/azimuth-mouse-lung-cellref-seed) lung on July 25^th^, 2023. The LungMAP consortium is funded by the National Heart, Lung and Blood Institute (NHLBI). Secondary annotation of cell identity was performed utilizing data generated by the Human Lung Cell Atlas version 2 utilizing the Azimuth interface (https://azimuth.hubmapconsortium.org/).

### Statistical analysis

Statistical comparisons between multiple groups were performed using one way ANOVA. Post hoc analyses were performed with Bonferroni’s corrections. Values are presented as the mean ± SEM and p < 0.05 was regarded as significant.

## ACKNOWLEDGMENTS

The authors would like to thank the laboratory of Lucy Golden, PhD in the Department of Medicine, Keck School of Medicine for assistance with FACS and scRNAseq library construction. Histology and microscopy services were provided by the Cell and Tissue Imaging Core of the University of Southern California (USC) Research Center for Liver Diseases (P30 DK048522 and S10 RR022508) and Norris Comprehensive Cancer Center Core (P30 CA014089). This work was supported by research grants R35 HL135747 (Z.B.) and R01 HL114959 (B.Z. & D.A.). CNM was supported by the National Cancer Institute (R01 CA 262258), the American Cancer Society through a Research Scholar Grant (RSG-20-135-01), the Wright Foundation, and the Departments of Surgery and of Translational Genomics at the Keck School of Medicine, USC. J.C. was supported by a Dean’s Fellowship from the Keck School of Medicine and the CaRE^2^ Florida-California Health Equity Center (U54 CA23396, U54 CA233444, and U54 CA233465).

## AUTHOR CONTRIBUTIONS

B.Z, C.N.M, and Z.B. supervised the work, designed all the experiments, and interpreted the data. H.S, P.F, B.Z. and Z.B generated the transgenic mouse model *Gramd2*^CreERT2^. H.S. maintained mouse strains and performed mouse experiments, 3-D culture and staining. W.C. and Y.L. performed 3-D culture and staining. A.C. processed cells for scRNAseq. Y.J maintained mouse strains. J.C., S.S., M.H., and C.N.M performed the bioinformatic analysis. C.L. and P.M. interpreted experimental data and edited the manuscript. H.S, C.N.M, B.Z, and Z.B. wrote the manuscript.

## INCLUSION AND DIVERSITY

One or more of the authors of this paper self-identifies as an underrepresented ethnic minority in science. One or more of the authors of this paper received support from a program designed to increase minority representation in science.

