## Supplemental Figures for "GRAMD2^+^ alveolar type I cell plasticity facilitates cell state transitions in organoid culture"

**SUPPLEMENTARY FIGURES**

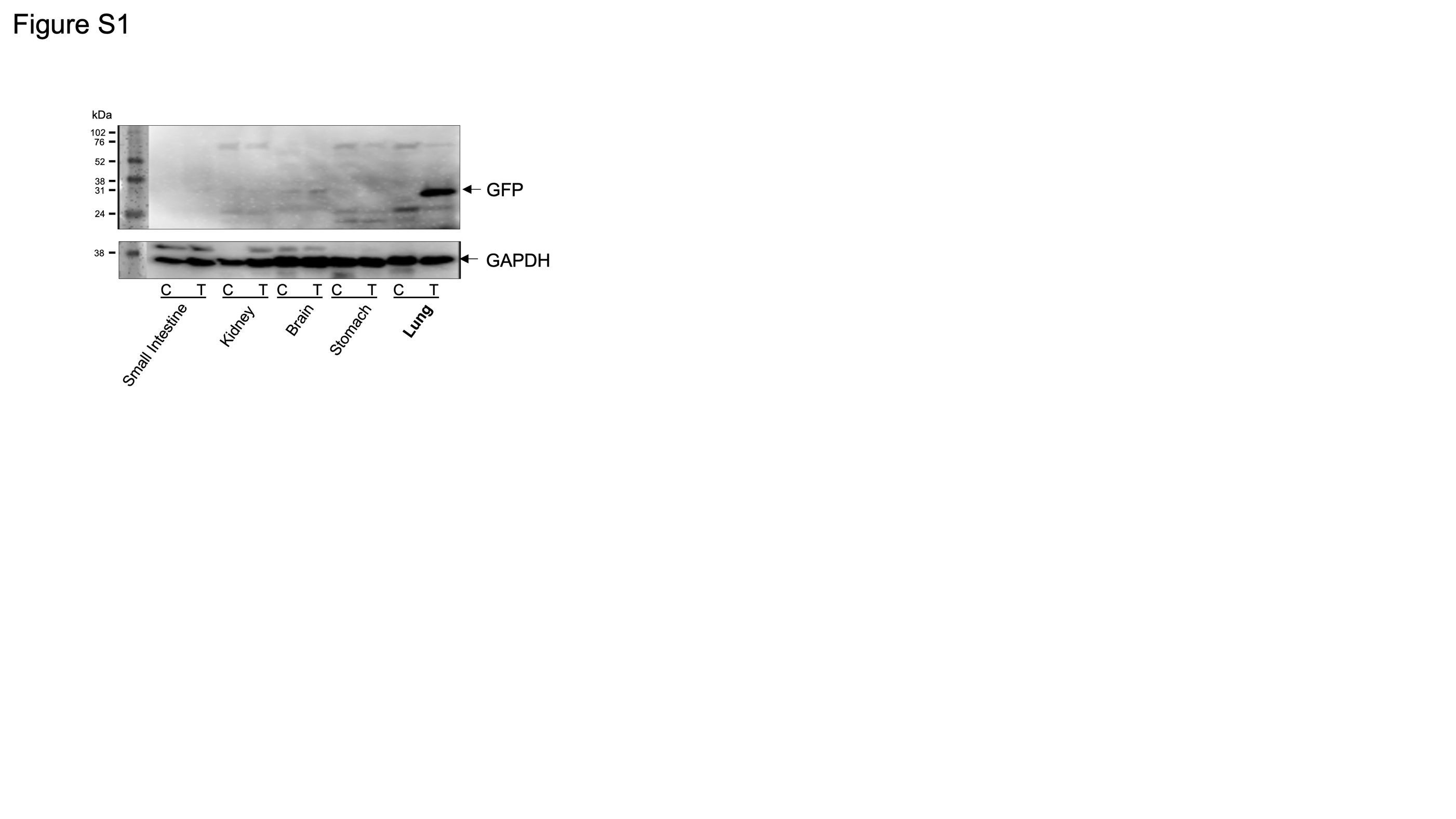

**Figure S1**. **Lineage labeling of lungs in** ***Gramd2^creERT2;mTmG^* mice.**

Representative western blot for GFP expression in *Gramd2^creERT2;mTmG^* mouse organs comparing tamoxifen-treated (T) and tamoxifen-untreated conditions (C). GAPDH is the loading control.

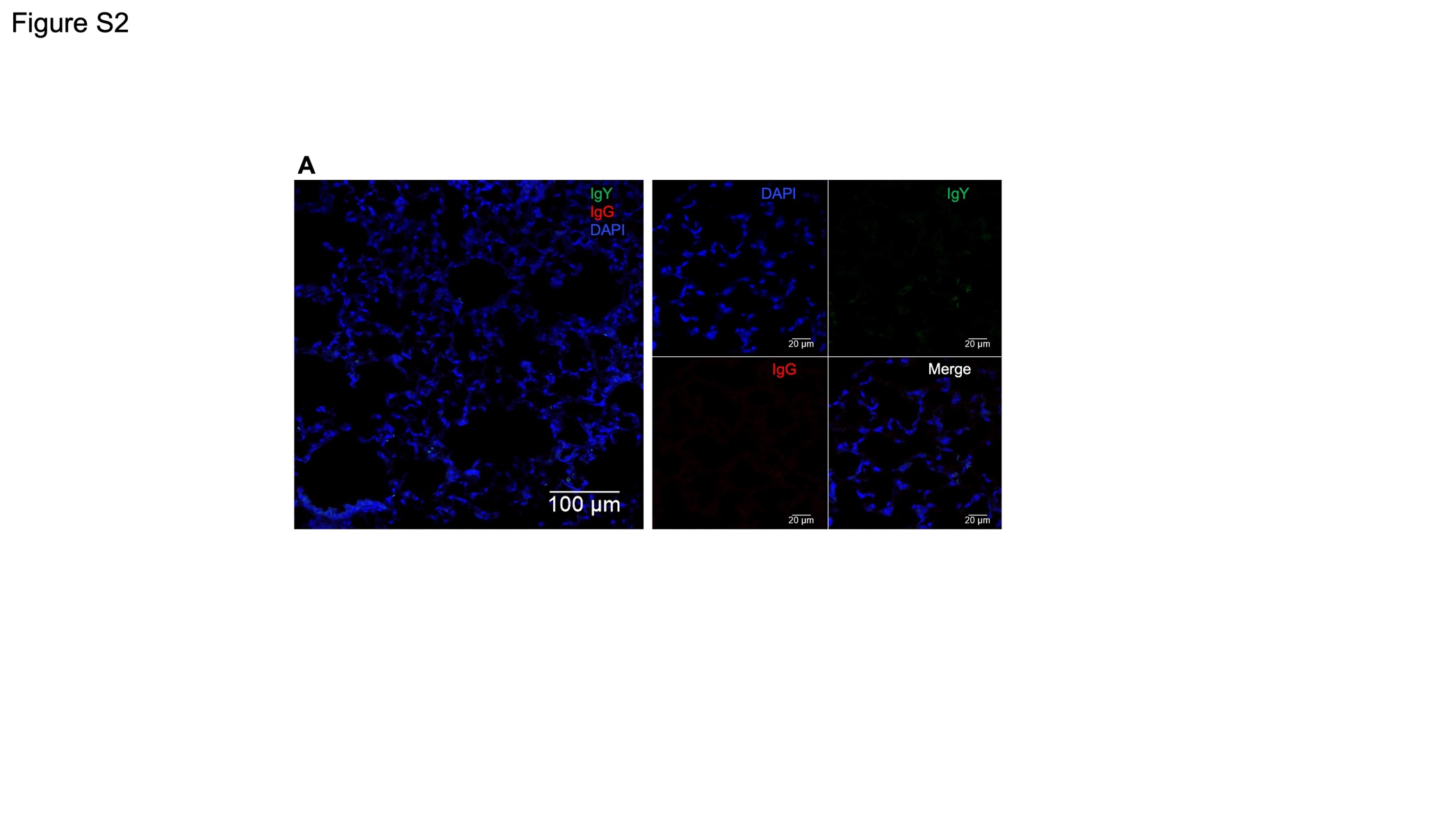

**Figure S2. Confocal images showing negative controls for GFP and AQP5) co-staining of *Gramd2^creERT2;mTmG^* lung sections.**

**A.** Representative confocal tile scan of chicken IgY (green) and rabbit IgG (red) in lungs of *Gramd2^creERT2;mTmG^* mice following tamoxifen (100 mg/kg) on two consecutive days. DAPI (blue) is the nuclear counterstain.

**B.** Confocal images of chicken IgY (green) and rabbit IgG (red). DAPI (blue) is the nuclear counterstain. N=3.

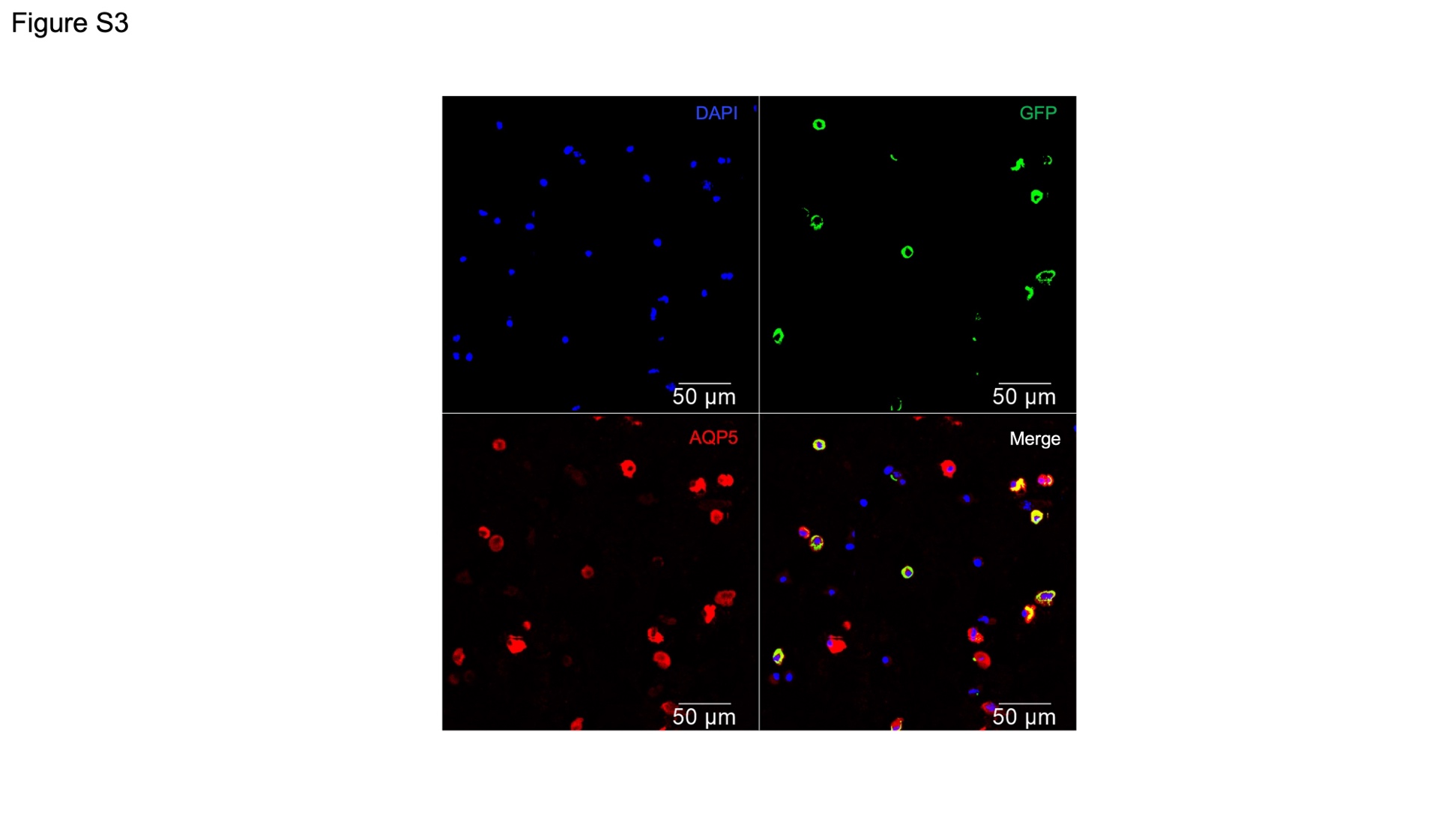

**Figure S3**. **Single channel images from Figure 1K**.

Representative cytospin staining of crude cell preparations from distal lung tissue of *Gramd2^creERT2;mTmG^* mice. Green = GFP, red = AQP5, blue = DAPI. N=3. Scale bar = 50 μM.

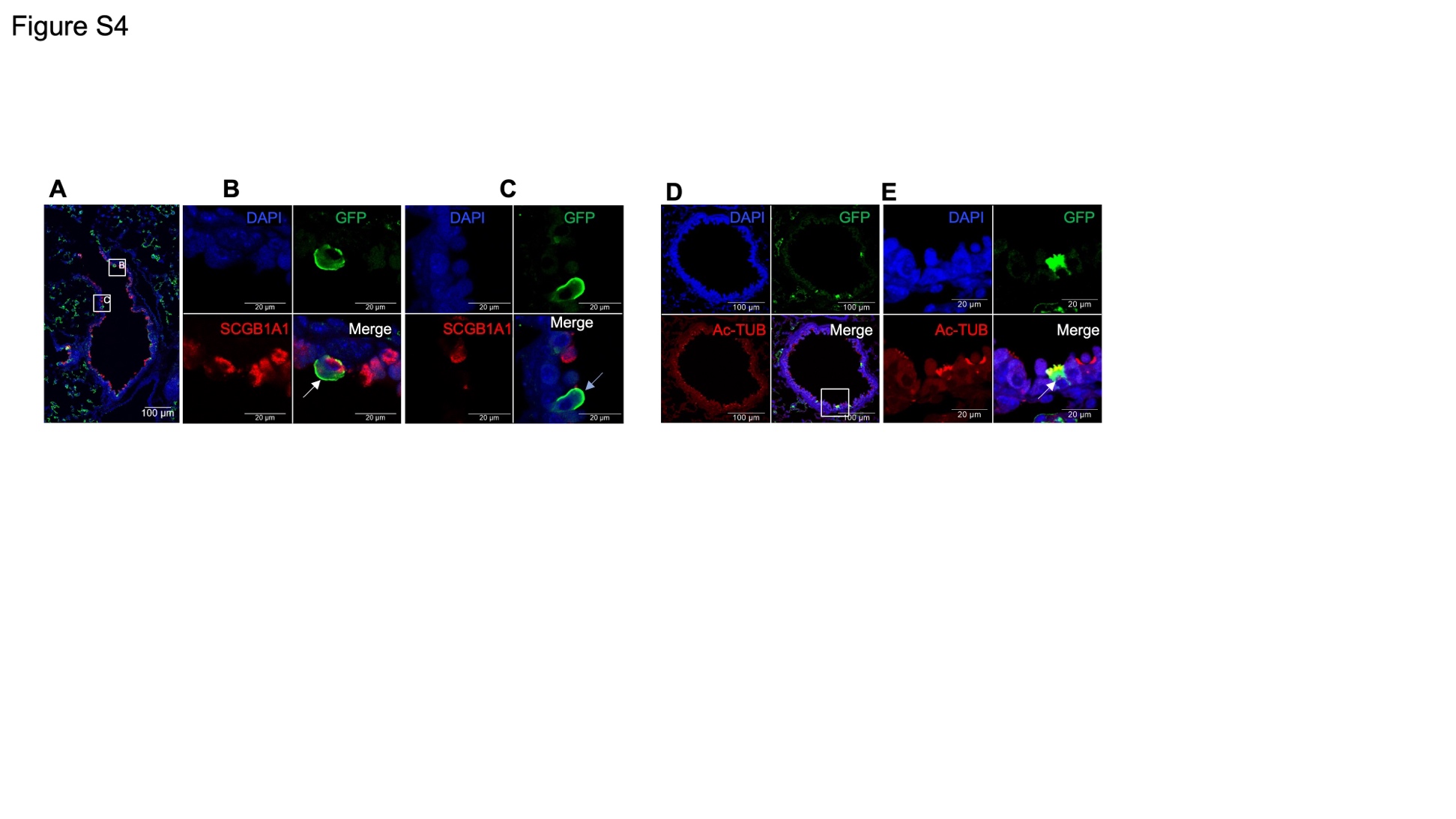

**Figure S4. Confocal images for club and ciliated cell staining of *Gramd2^creERT2;mTmG^* lung sections.**

**A.** Low-magnification confocal image of GFP (green), SCGB1A1 (red), and DAPI (blue) in *Gramd2^creERT2;mTmG^* paraformaldehyde-fixed, paraffin - embedded lung sections. Scale bar (white) = 100 μm.

**B**. Higher-magnification confocal images of *Gramd2^creERT2;mTmG^* paraformaldehyde-fixed, paraffin - embedded lung sections centered on the upper white box in (A). Cells are colored by GFP (green), SCGB1A1 (red), and DAPI (blue) Individual and merged 3-color channels are shown. The white arrow indicates an identified dual-positive GFP^+^SCGB1A1^+^ cell. Scale bar (white) = 20 μm.

**C.** Higher-magnification confocal images of *Gramd2^creERT2;mTmG^* paraformaldehyde-fixed, paraffin - embedded lung sections centered on the lower white box in (A). Cells are colored by GFP (green), S CGB1A1 (red), and DAPI (blue). Individual and merged 3-color channels are shown. The cerulean arrow indicates an identified dual-positive GFP^+^SCGB1A1^-^ cell. Scale bar (white) = 20 μm.

**D**. Low-magnification confocal imaging of GFP (green), acetylated tubulin (Ac-TUB, red), and DAPI (blue) of *Gramd2^creERT2;mTmG^* paraformaldehyde-fixed, paraffin - embedded lung sections. Individual and merged 3-color channels are shown. Scale bar (white) = 100 μm.

**E**. Higher-magnification confocal images of*Gramd2^creERT2;mTmG^* paraformaldehyde-fixed, paraffin - embedded lung sections centered on the white box in (D). Staining for GFP (green), acetylated tubulin (Ac-TUB), and DAPI (blue). Individual and merged 3-color channels are shown. The white arrow indicates a dual-positive GFP^+^ Ac-TUB^+^ cell. Scale bar (white) = 20 μm.

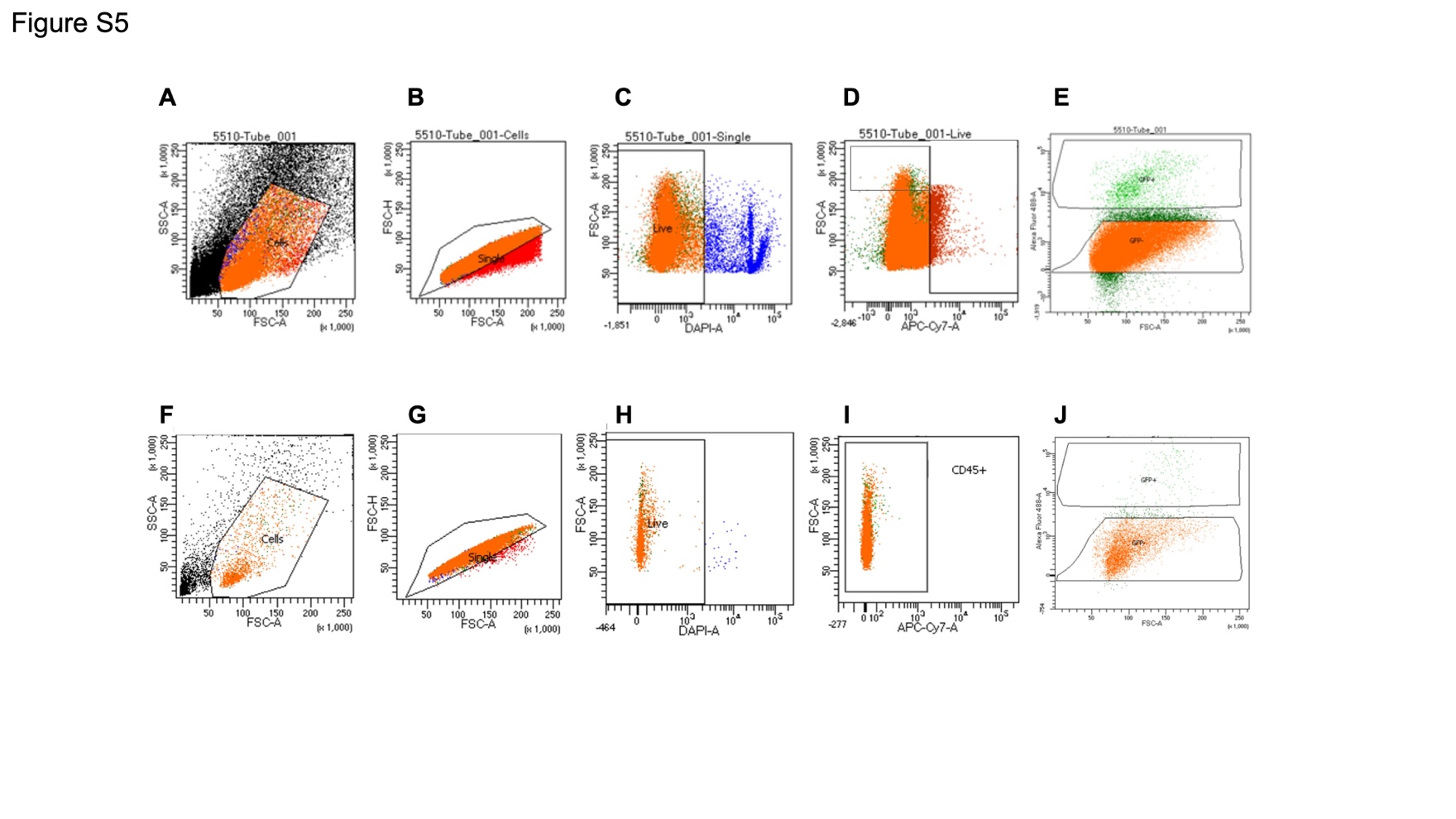

**Figure S5. Gating strategy to identify GFP^+^ AT1 cells from *Gramd2^CreERT2;mTmG^* mice** **by FACS.**

**A.** Tamoxifen-treated *Gramd2^CreERT2;mTmG^* single cell suspensions from distal lung isolates were gated based on forward scatter (FSC) versus side scatter (SSC) to remove debris. N=8. Representative sorting shown.

**B.** *Gramd2^CreERT2;mTmG^* cells filtered in (**A**) then had doublets removed via area the cell occupies (FSC-A) versus the height of the cell (FSC-H). N=8. Representative sorting shown.

**C.** *Gramd2^CreERT2;mTmG^* cells remaining from (**B**) then were filtered based on DAPI staining; those with high levels of DAPI were excluded. N=8. Representative sorting shown.

**D.** *Gramd2^CreERT2;mTmG^* cells remaining from (**C**) were then filtered to remove any contaminating immune cells by positive selection of cells with little to no CD45 (ApC-Cy7-A) staining. N=8. Representative sorting shown.

**E.** Lastly, cells from (**D**) were separated based on intensity of GFP signal using Alexa Flour 488-A levels. N=8. Representative sorting shown.

**F.** GFP antibody-treated wild type (129SvEv/Tac) distal lung single cell suspension isolates, which lack the *Gramd2^CreERT2;mTmG^* transgene, were used to determine the GFP^-^ population for flow cytometry gating. Initial gating was performed as in (**A**). N=8.

**G.** Cells selected for in (**F**) then had doublets removed via area the cell occupies (FSC-A) versus the height of the cell (FSC-H). N=8. Representative sorting shown.

**H.** Cell suspensions filtered for in (**G**) were subsequently filtered based on DAPI staining; those with high levels of DAPI were excluded as in (**C**). N=8. Representative sorting shown.

**I.** Cells selected for in (**H**) then had any contaminating immune cells depleted by selecting cells with little to no CD45 (ApC-Cy7-A) staining as in (**D**). N=8. Representative sorting shown.

**J.** Lastly, cells from (**I**) were gated to separate GFP^+^ and GFP- populations based on Alexa Flour 488-A levels, as was performed in (**E**). N=8. Representative sorting shown.

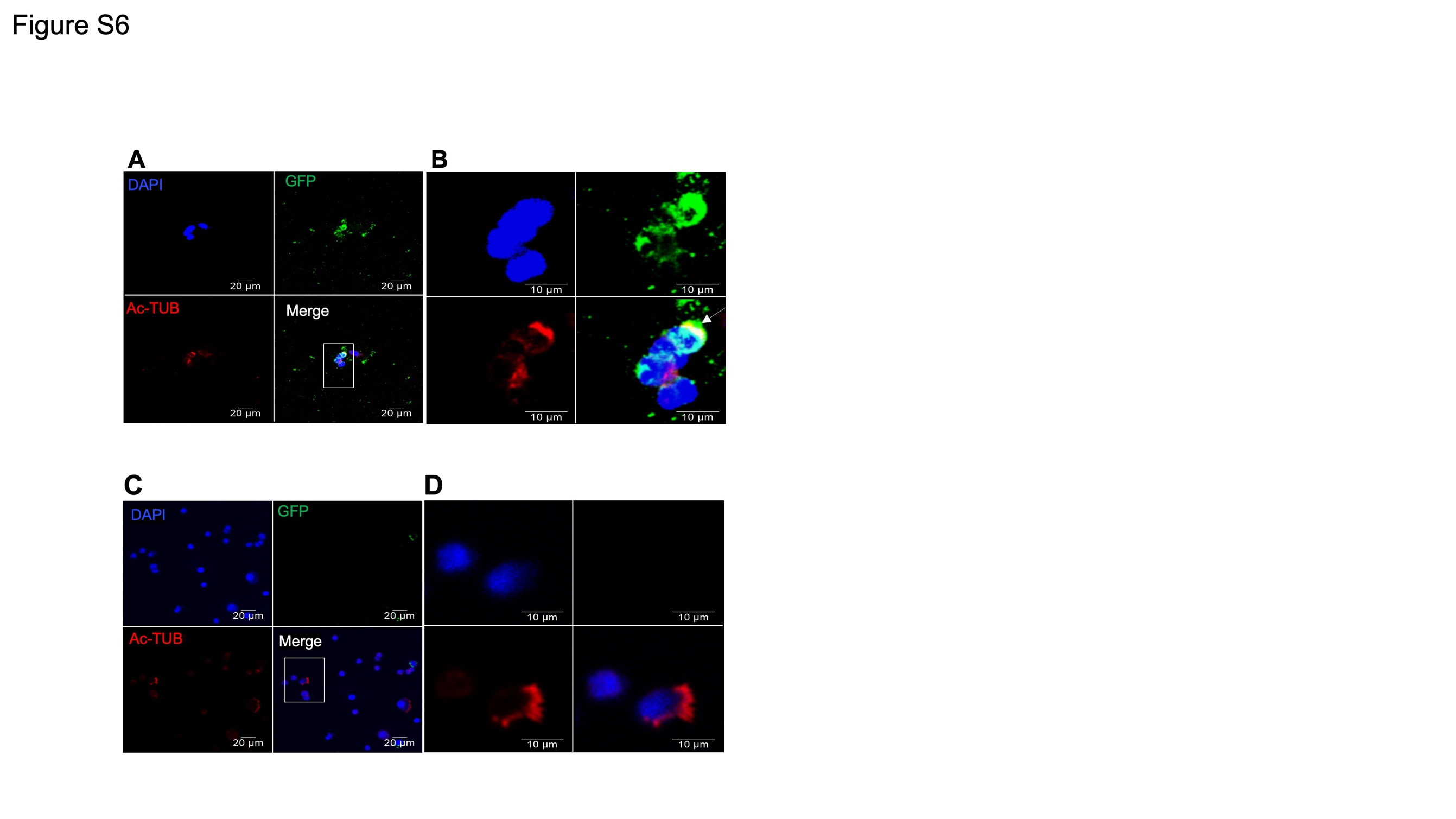

**Figure S6. Double staining for GFP and Ac-Tub in FACS cytospins.**

**A**. Cytospin staining of FACS sorted *Gramd2^CreERT2;mTmG^* GFP^+^ cells. GFP (green) was visualized together with staining for acetylated tubulin (Ac-TUB), a marker of ciliated cells. Individual and merged 3-color channels are shown. Scale bar (white) = 20 μm.

**B**. Cytospin staining of FACS sorted GFP^+^ cells. GFP (green) was visualized together with staining for acetylated tubulin (Ac-TUB). Individual and merged 3-color channels are shown. Scale bar (white) = 10 μm.

**C.** Cytospin staining of FACS sorted *Gramd2^CreERT2;mTmG^* GFP- cells. GFP (green) was visualized together with staining for acetylated tubulin (Ac-TUB). Individual and merged 3-color channels are shown. Scale bar (white) = 20 μm.

**D**. Cytospin staining of FACS sorted GFP- cells. GFP (green) was visualized together with for acetylated tubulin (Ac-TUB). Individual and merged 3-color channels are shown. Scale bar (white) = 10 μm.

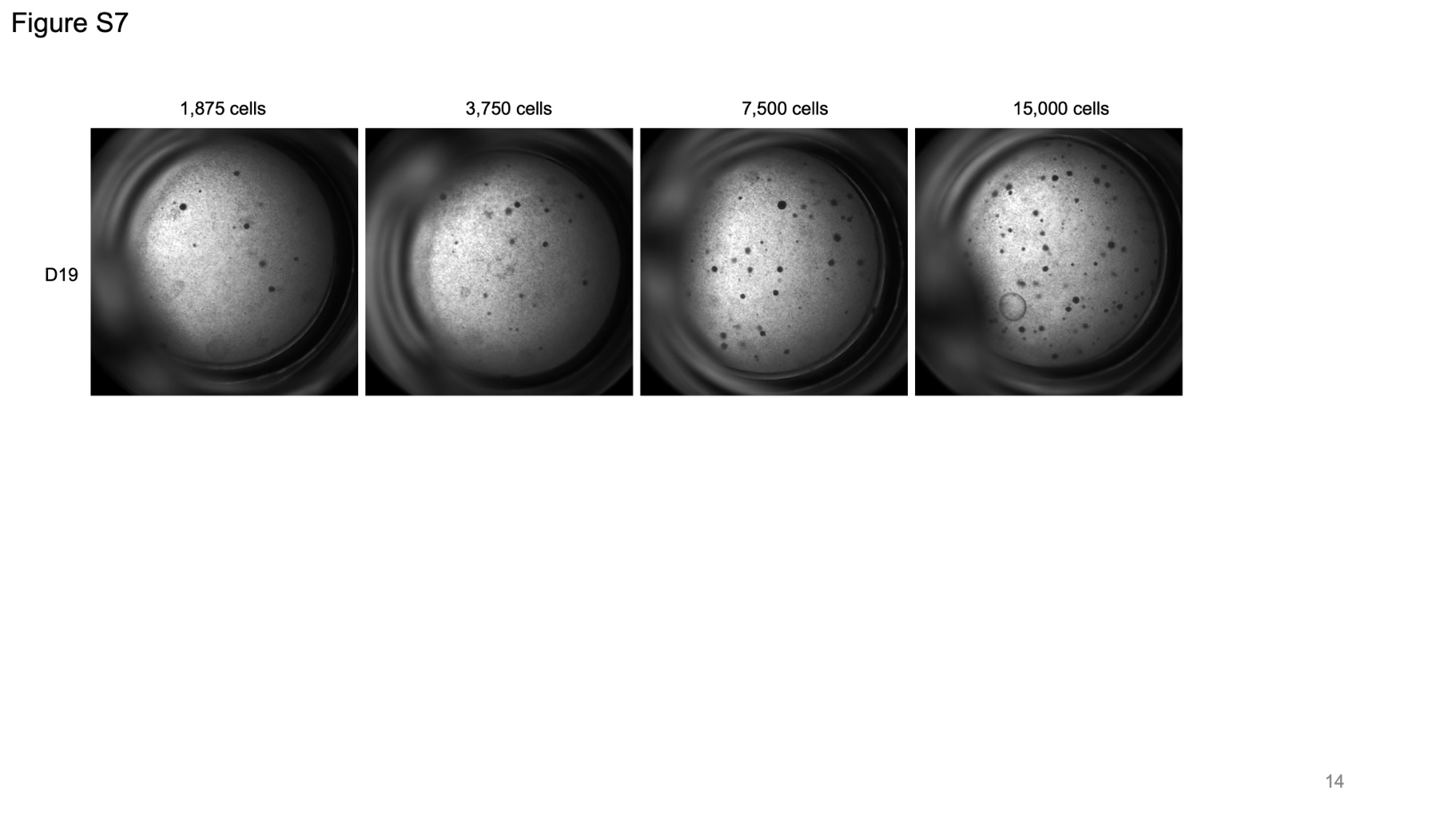

**Figure S7. Brightfield images of organoids formed from GFP^+^ cells from *Gramd2^CreERT2;mTmG^* mice.**

Number of colonies increases with increased plating density, but CFE remains the same.

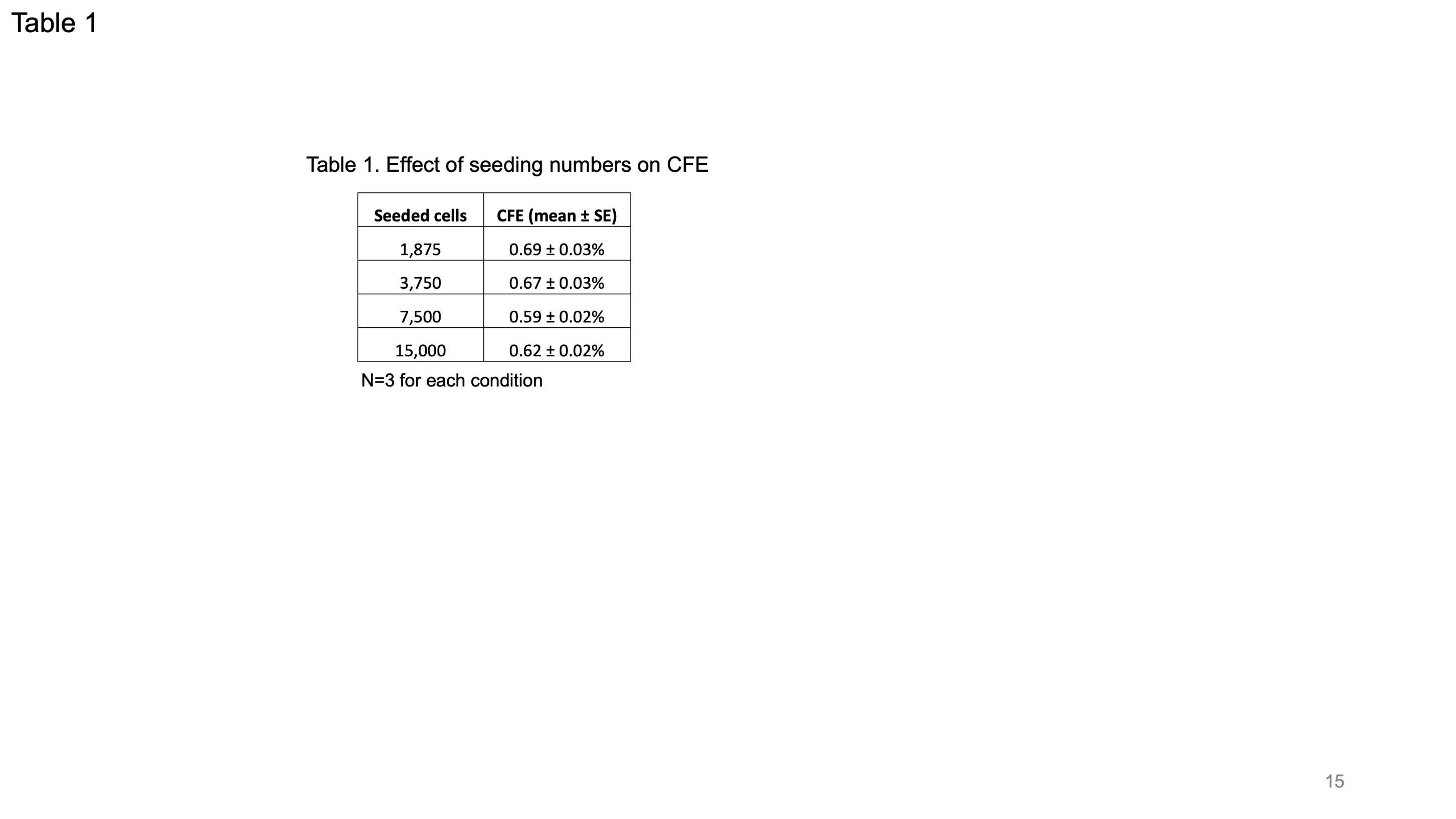

**Table S1. Effect of seeding density on organoid forming efficiency (CFE)**. N=3.

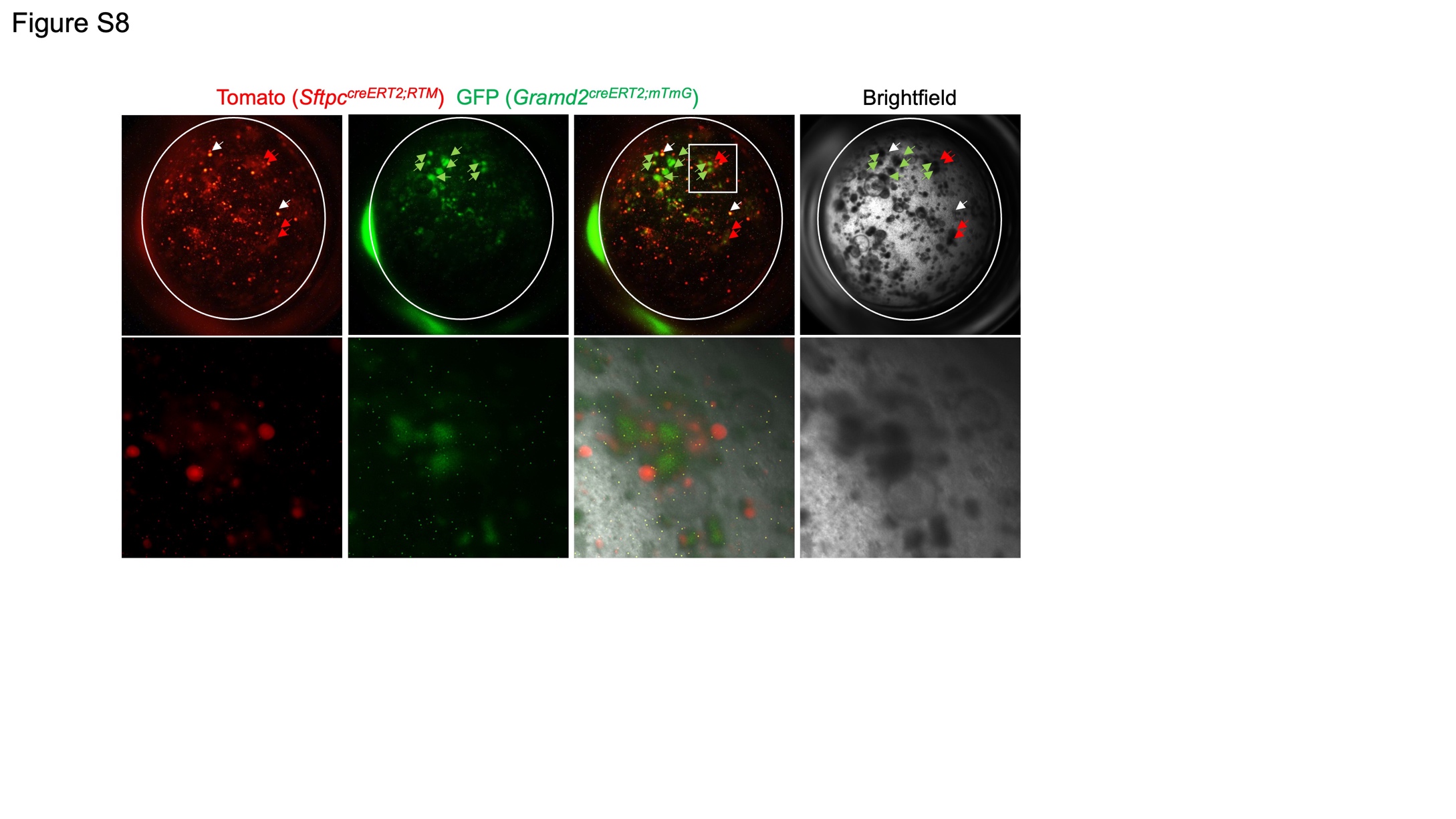

**Figure S8. Representative images show that AT1 and AT2 cells do not form mixed organoids.**

Cells from *Sftpc^creERT2;RTM^* mouse lungs were processed for AT2 cell preparation and RTM^+^ cells were collected by flow cytometry. Cells from *Gramd2^creERT2;mTmG^* mouse lung were processed for AT1 cell preparation and GFP^+^ cells were collected by flow cytometry. 10,000 RTM^+^ and 10,000 GFP^+^ cells were co-cultured with Mlg fibroblasts in Matrigel. Images show that on day 16, organoids were either RTM^+^ (AT2 cells, red arrow) or GFP^+^ (AT1 cells, green arrow). The yellowish dots (white arrow) are overexposed to see week RTM. Lower panel is higher magnification of boxed area in upper panel.

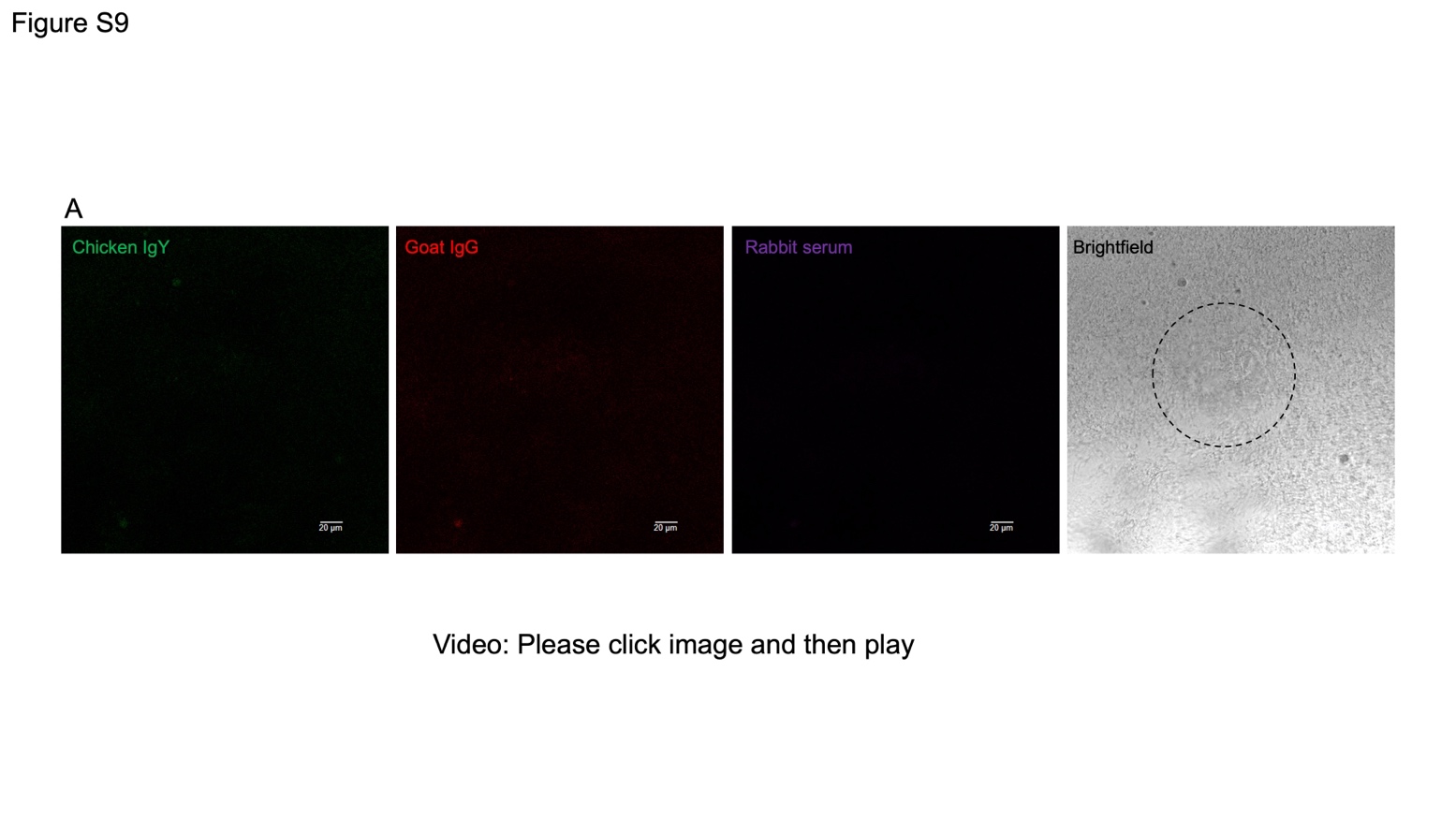

**Figure S9. Negative control for organoid whole mount IF.**

Exported video of confocal Z-stack images of whole mount IF for organoids on day 25 with individual channels (chicken IgY (green), goat IgG (red), rabbit serum (far-red channels) and bright field Circled area is the organoids. N=3.

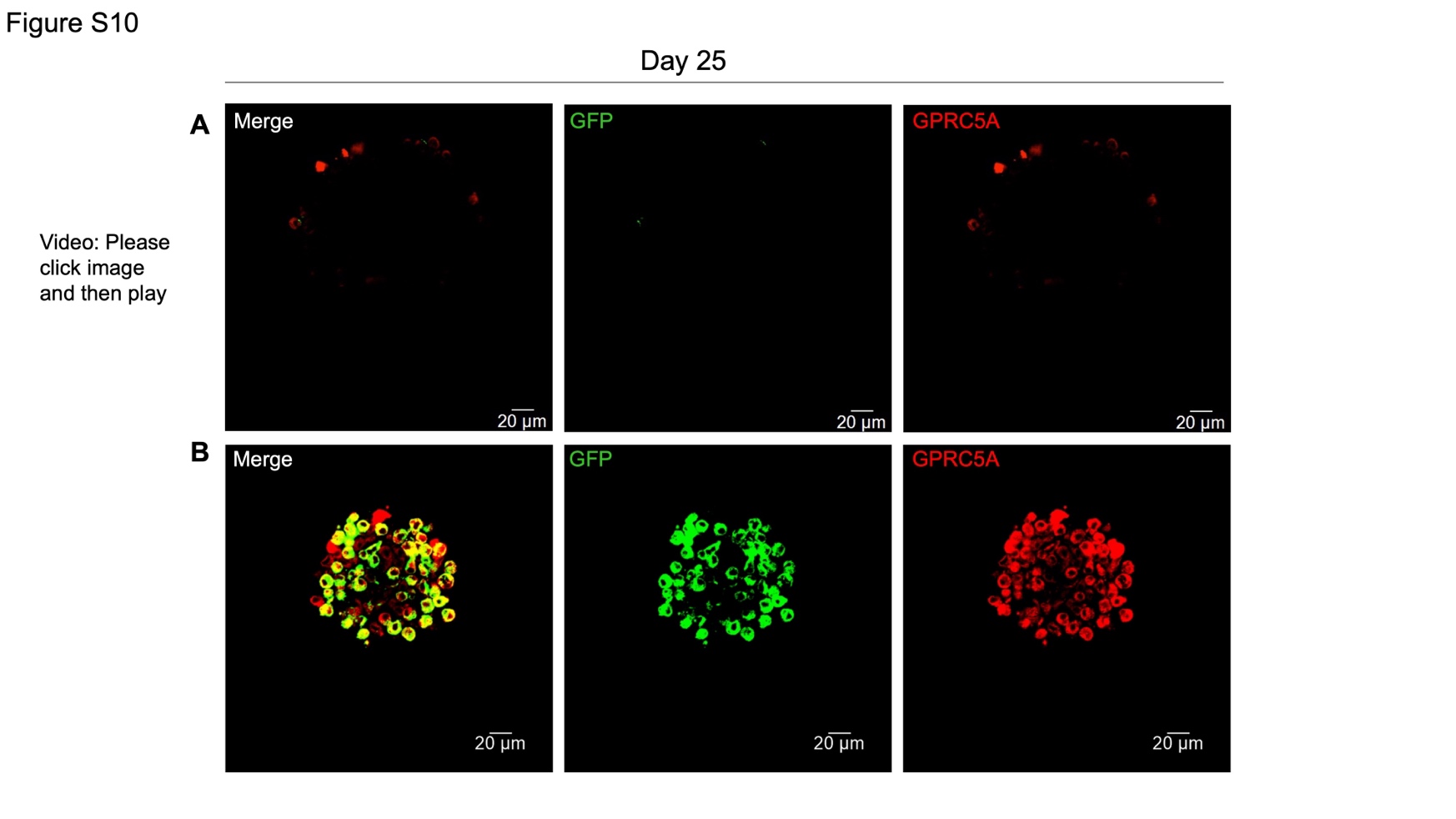

**Figure S10. Whole mount IF shows that GFP colocalizes with GPRC5A in organoids.**

**A.** Exported video (A) of confocal Z-stack images. Scale bar (white) = 20 μm.

**B**. representative confocal images of merged and single channels. GFP (green) and GPRC5A (red) for whole mount IF. Scale bar (white) = 20 μm.

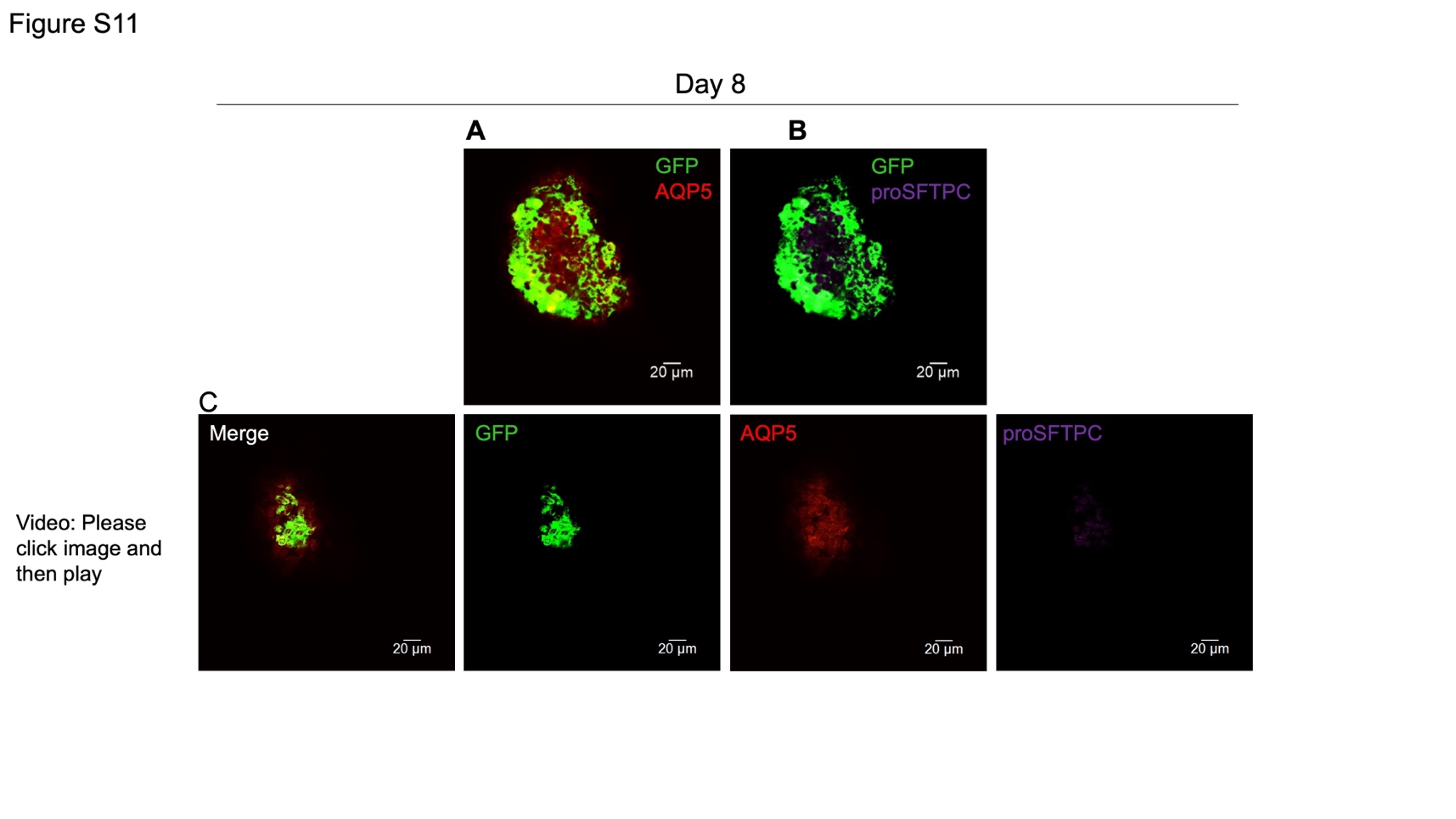

**Figure S11. Representative organoid of GFP^+^ cells from *Gramd2^creERT2;mTmG^* mice** **on day 8.**

**A.** Confocal image of GFP (green) costained with AQP5 (red) . Merged color channels are shown. Scale bar (white) = 20 μm.

**B.** Confocal image of GFP (green) costained with proSFTPC (purple). Merged color channels are shown. Scale bar (white) = 20 μm.

**C.** Confocal Z-stack video of an organoid that underwent whole mount IF imaging on day 8. GFP (green), AQP5 (red) and proSFTPC (purple) were co-stained. Merged color channels are shown. Scale bar (white) = 20 μm.

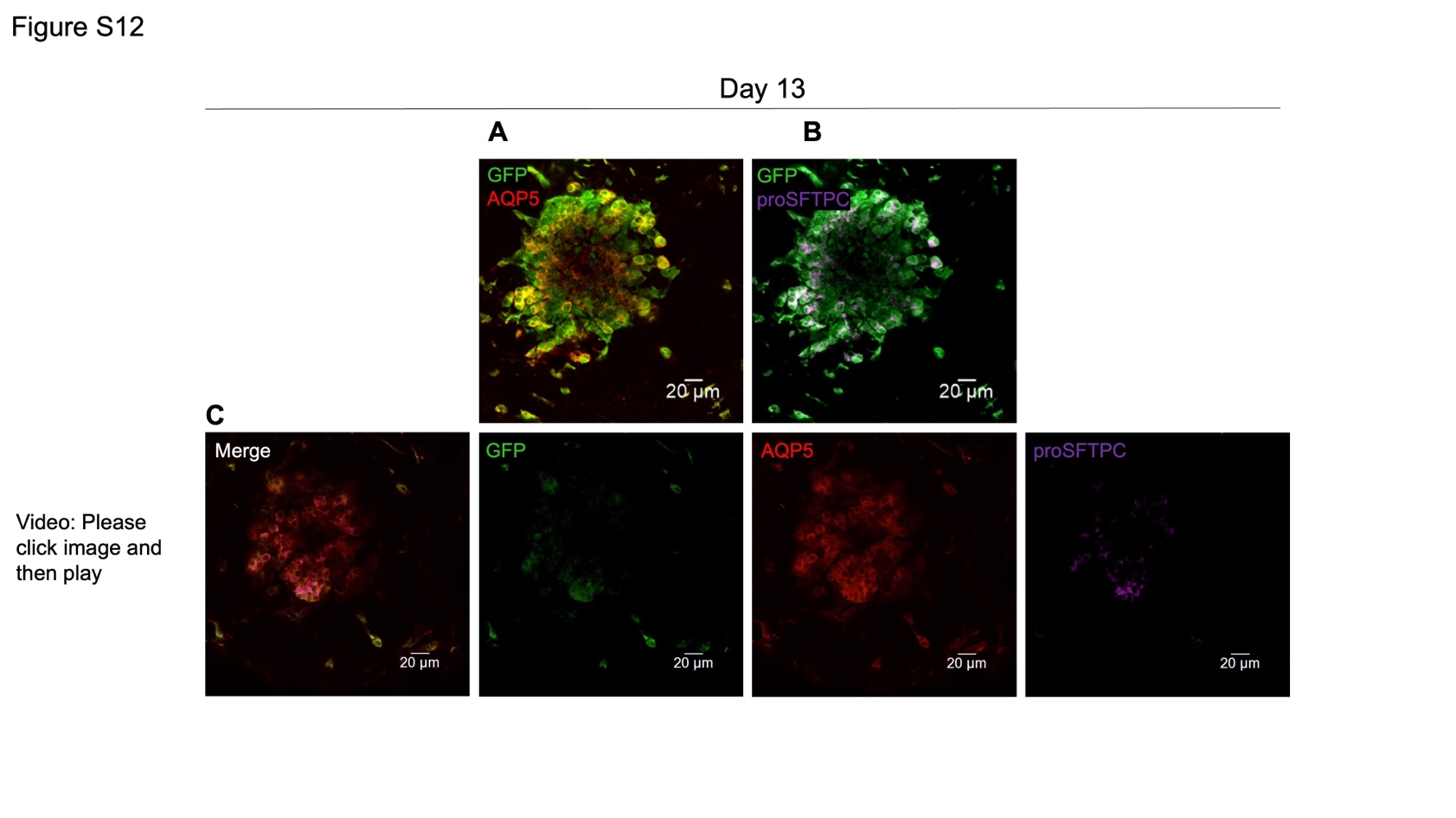

**Figure S12. Representative organoid of GFP^+^ cells from *Gramd2^creERT2;mTmG^* mice** **on day 13.**

**A.** Confocal image of GFP (green) costained with AQP5 (red). Merged color channels are shown. Scale bar (white) = 20 μm.

**B.** Confocal image of GFP (green) costained with proSFTPC (purple). Merged color channels are shown. Scale bar (white) = 20 μm.

**C.** Confocal Z-stack video of an organoid that underwent whole mount IF imaging on day 13. GFP (green), AQP5 (red) and proSFTPC (purple) were co-stained. Merged color channels are shown. Scale bar (white) = 20 μm.

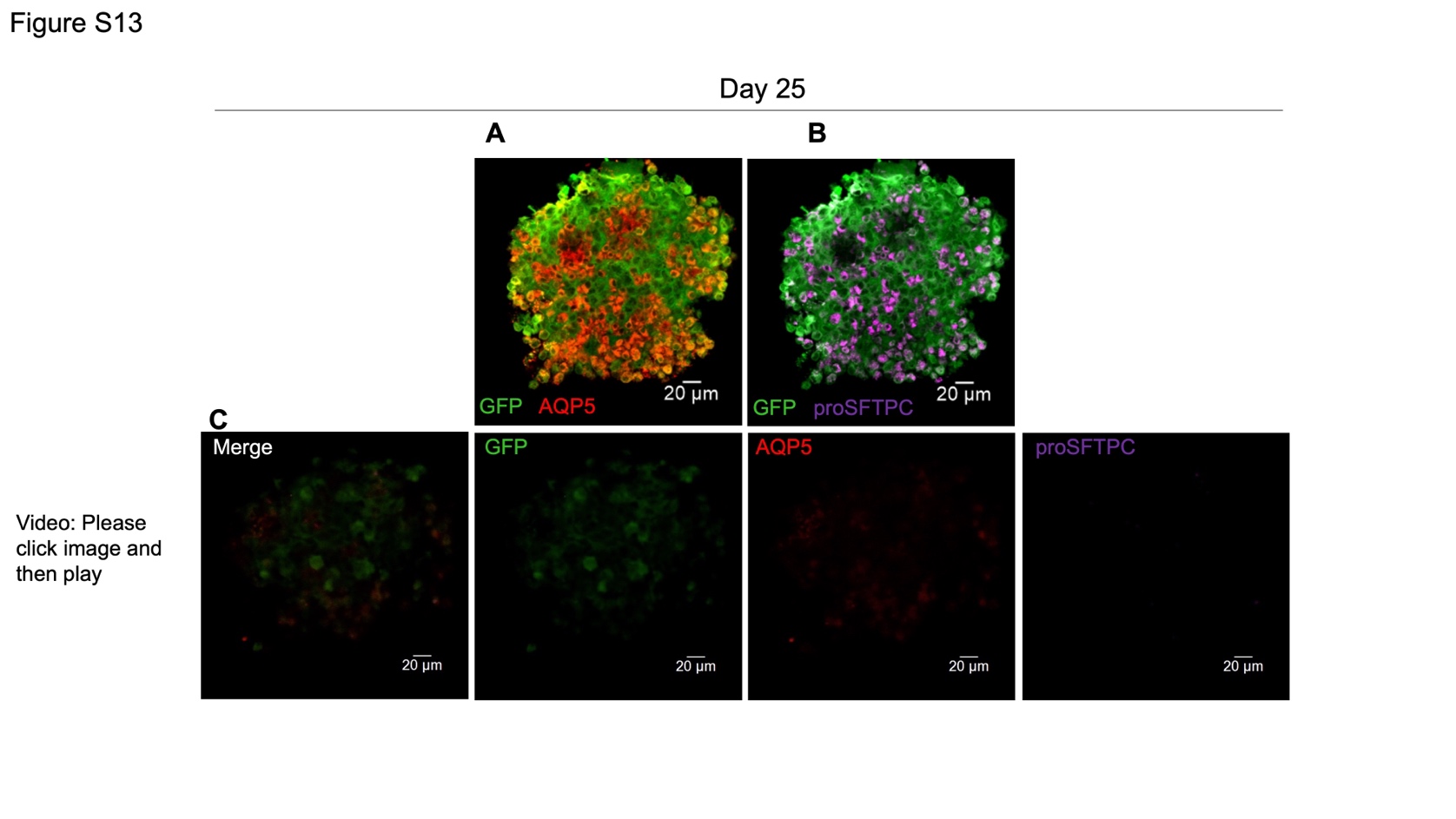

**Figure S13. Representative organoid of GFP^+^ cells from *Gramd2^creERT2;mTmG^* mice** **on day 25.**

**A.** Confocal image of GFP (green) costained with AQP5 (red). Merged color channels are shown. Scale bar (white) = 20 μm.

**B.** Confocal image of GFP (green) costained with proSFTPC (purple). Merged color channels are shown. Scale bar (white) = 20 μm.

**C.** Confocal Z-stack video of an organoid that underwent whole mount IF imaging on day 25. GFP (green), AQP5 (red) and proSFTPC (purple) were co-stained. Merged color channels are shown. Scale bar (white) = 20 μm.

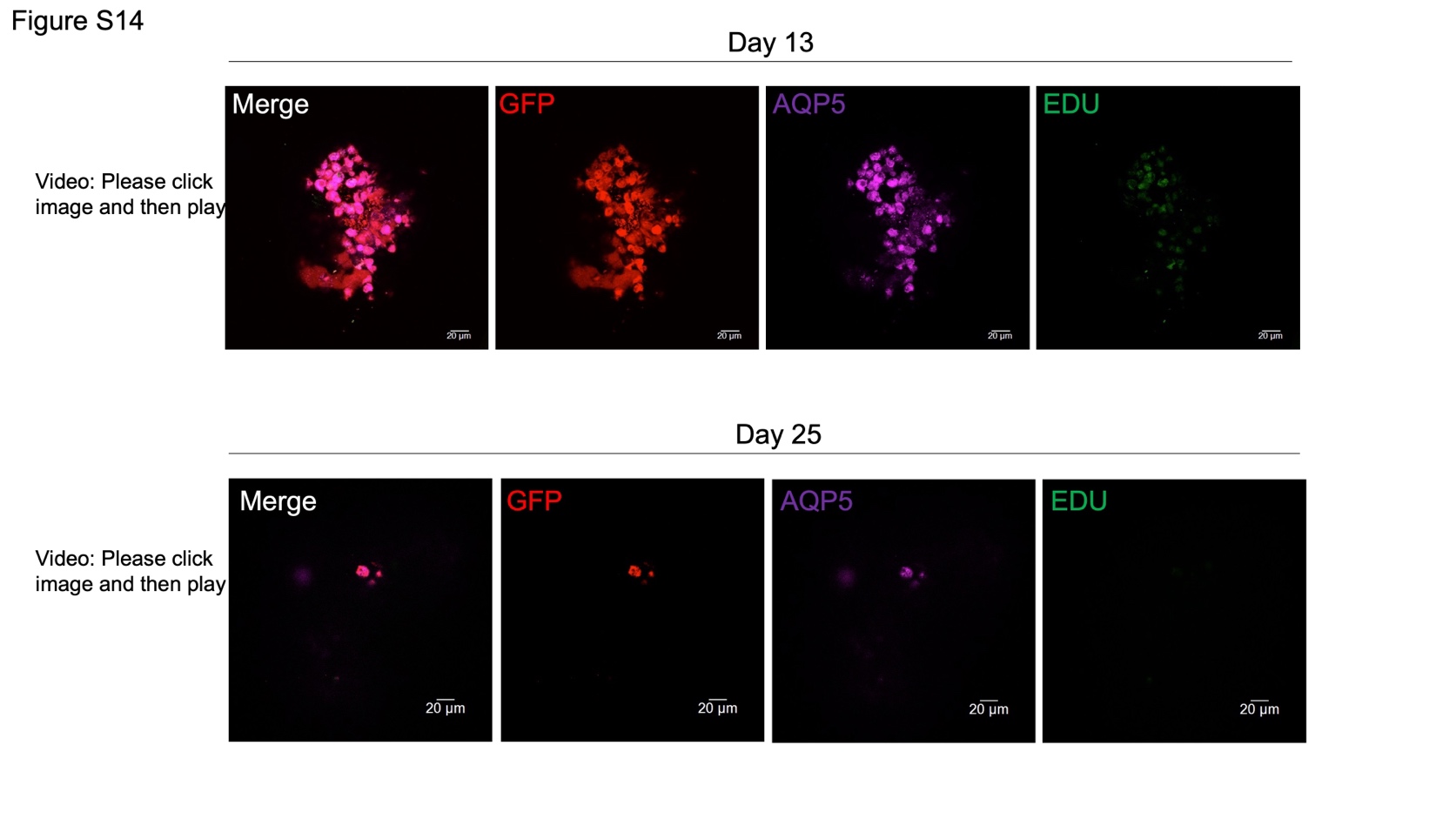

**Figure S14. Some GFP^+^AQP5^+^ cells are labeled by EdU in** **organoid culture on days 13 and 25.**

Exported video of confocal Z-stack images for GFP (red), AQP5 (purple), and EDU (green) of whole mount IF showing merged and single channels. Scale bar (white) = 20 μm.

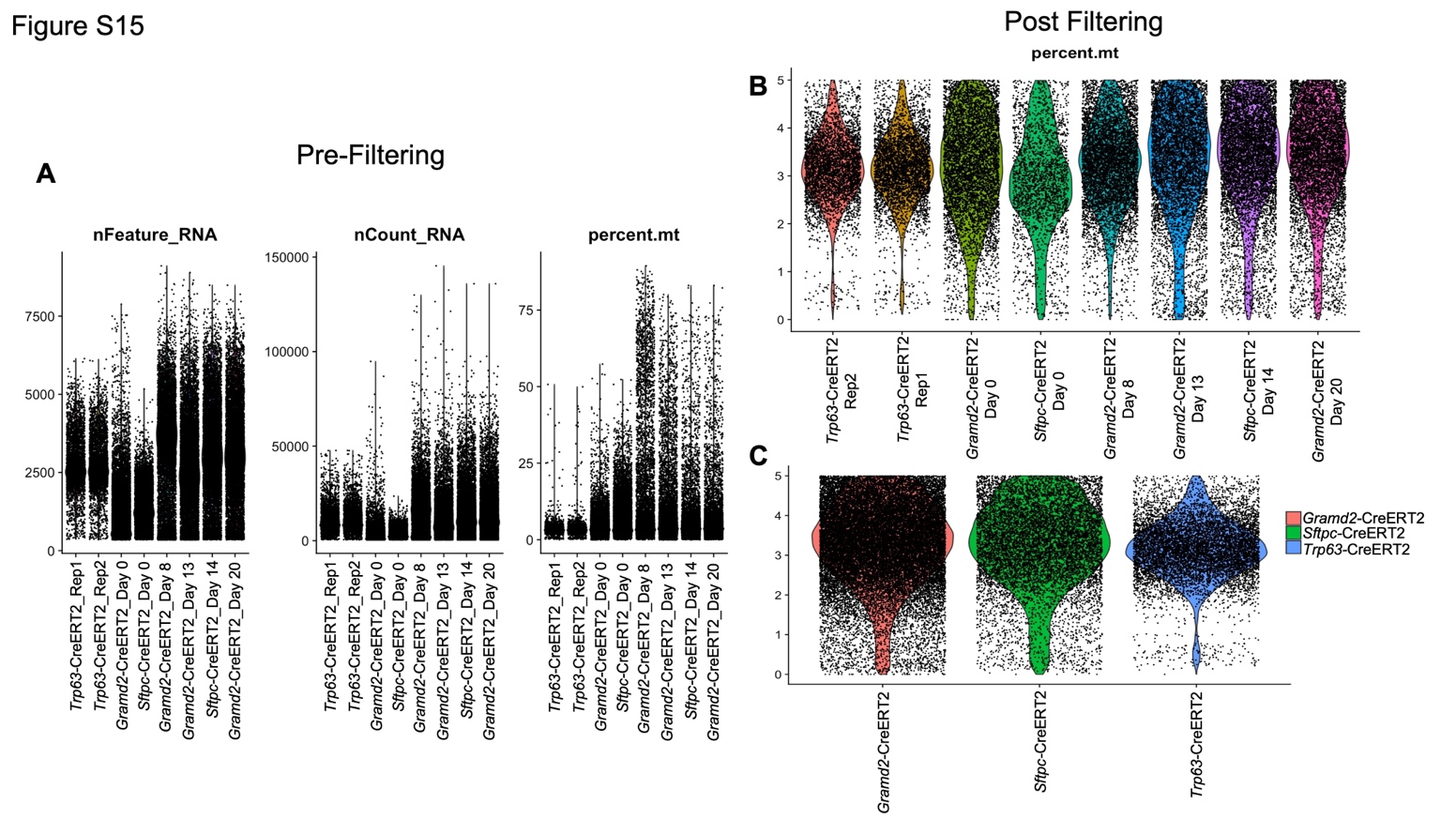

**Figure S15. Quality control metrics for scRNAseq analysis**.

**A.** Distribution of total genes expressed per cell (n_Feature), total number of reads aligned to exons (n_Counts), and percent mitochondrial contamination as a measure of cell viability (percent.mt) per cell in each sample. Distributions of *Gramd2^creERT2;mTmG^* samples (D0, D8, D13, and D20) were compared to previously published *Sftpc^CreERT2;TdTomato^* primary (D0) uncultured and organoid cultured D14 (D14), as well as primary uncultured *Trp63^CreERT2^* samples (D0 - Rep1, D0 - Rep2).

**B.** Violin distribution of cells based on percent mitochondrial levels post-filtering for percent.mt to retain only cells with less than 5% contamination (<5% retained). **C**. Violin distribution of cells by quality metrics aggregated by cell of origin. Red = *Gramd2^CreERT2^,* green= *Sftpc^CreERT2^*, and blue = *Trp63*^CreERT2^.

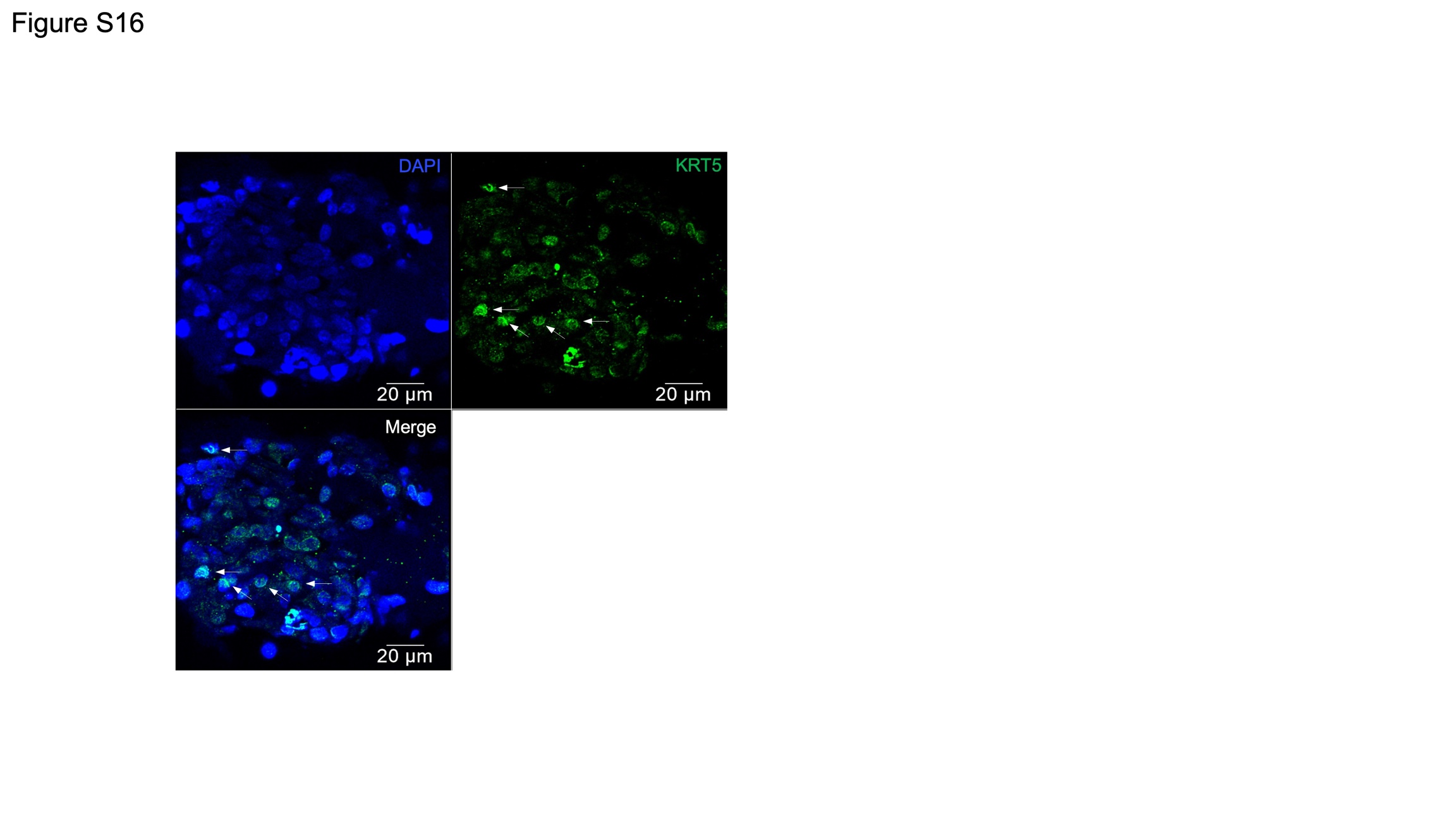

**Figure S16. KRT5 staining in organoids.**

Confocal IF image of organoid stained with KRT5 (green) in paraffin-embedded organoid sections on day 13. DAPI is nuclear counterstain. White arrow: KRT5^+^ cells. Scale bar (white) = 20 μm.

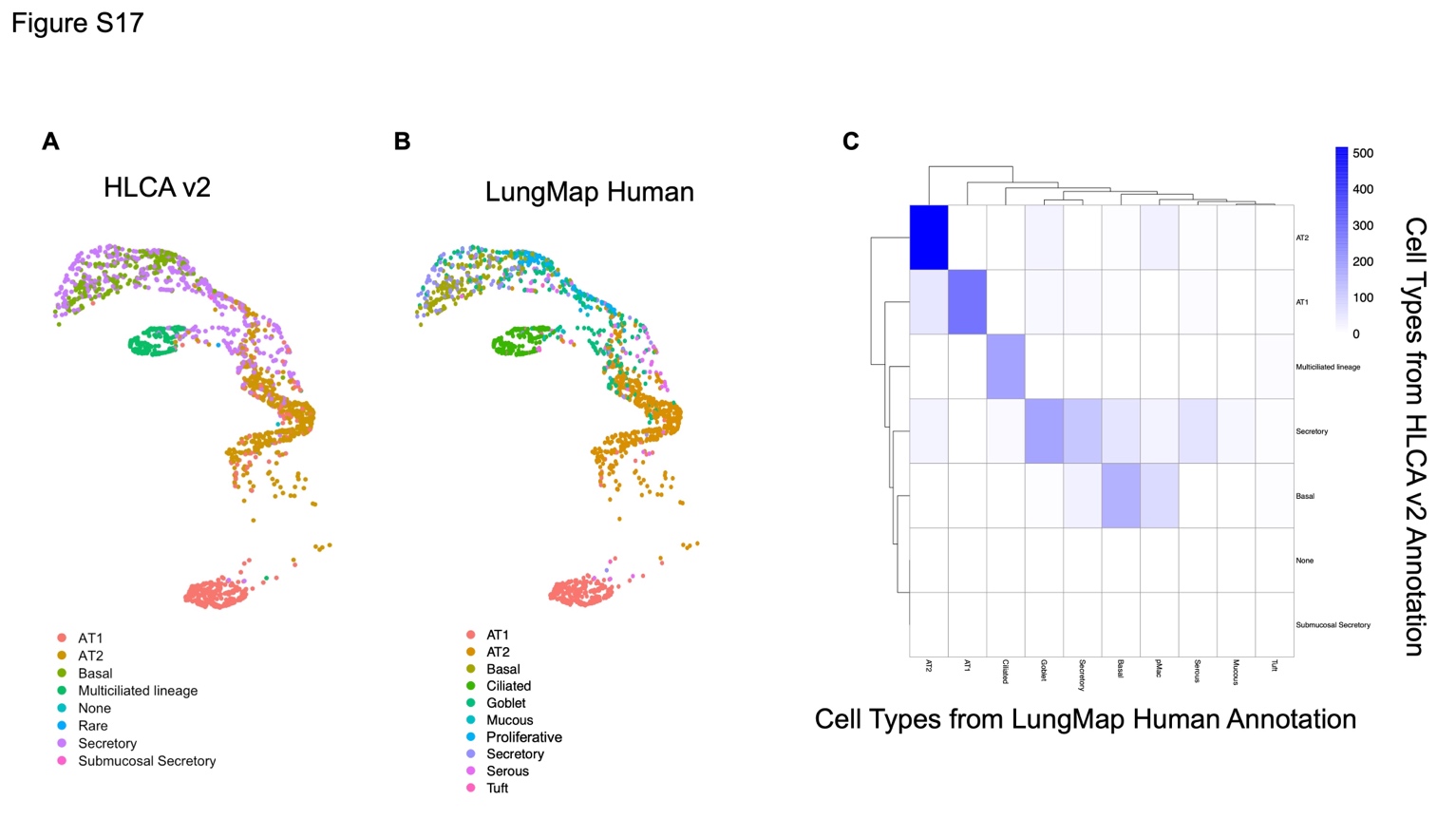

**Figure S17. Comparison of scRNAseq annotation between HLCAv2 and LungMAP**.

**A.** UMAP of EpCAM^+^ epithelial cell population annotated to human lung cell atlas version 2 (HLCA v2) using Azimuth. Red = AT1, gold = AT2, peridot = basal, green = multiciliated, teal = none, blue = rare, purple = secretory, pink = submucosal secretory.

**B.** UMAP of EpCAM^+^ epithelial cell population annotated to human LungMAP using Azimuth. Red = AT1, gold = AT2, peridot = basal, green = ciliated, teal = goblet, seafoam = mucous, blue = proliferative, purple = secretory, maroon = serous, pink = tuft.

**C**. Comparison of cell identity assignment between human LungMAP (x-axis) and HLCAv2 (y-axis). Blue = high correlation between annotations, white = low correlation between reference annotations.

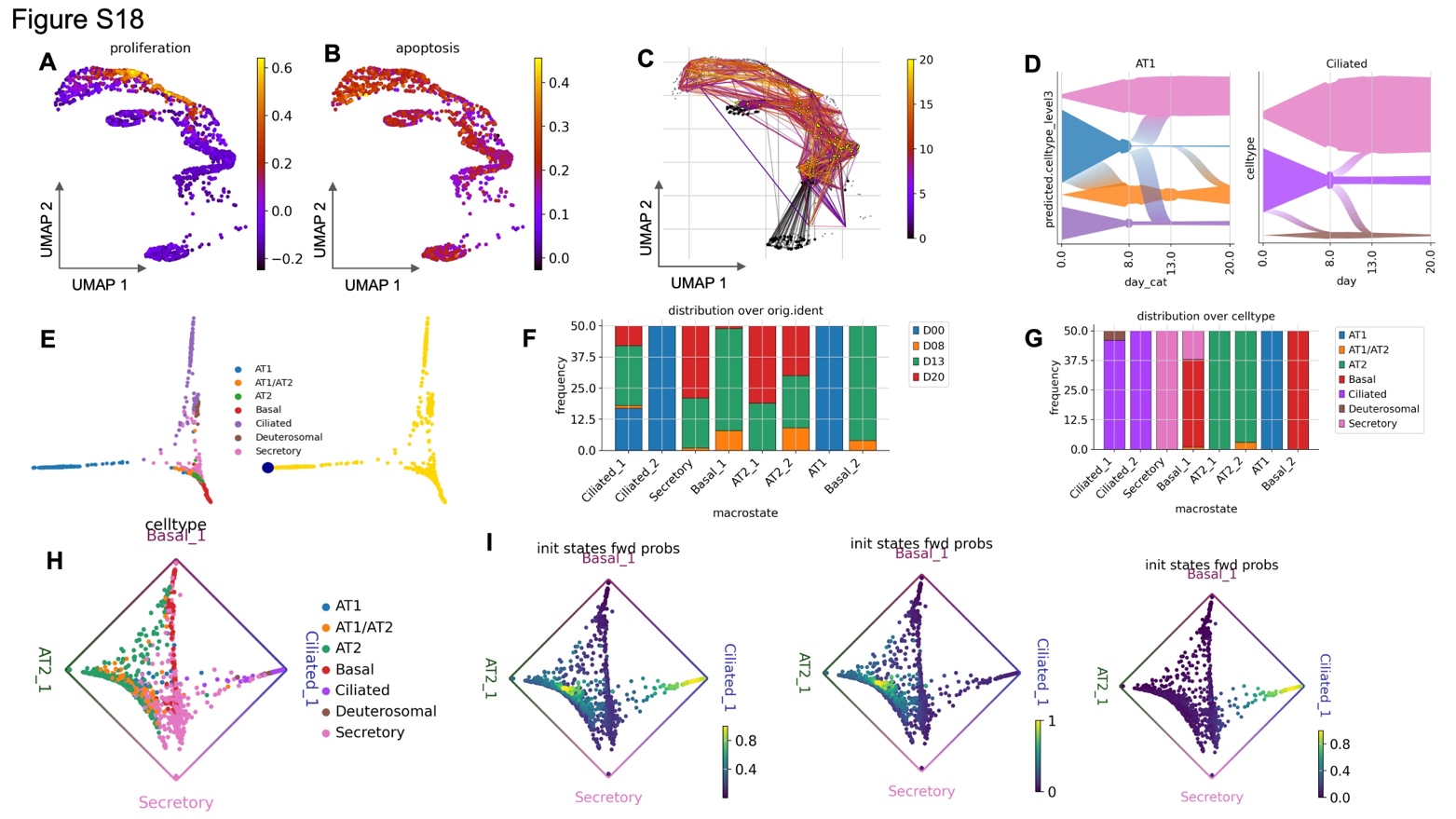

**Figure S18**. **CellRank transition matrix and estimator.**

**A-B.** To reconstruct differentiation trajectories across time points, moscot (multi-omic single-cell optimal transport tools) (43) was used to couple cells across time points using optimal transport (OT). Both proliferation and apoptosis gene signature scores were used to guide coupling across time points.

**C.** Psuedotime was measured using diffusion pseudotime (DPT) (44). The root cell used for computing DPT is shown on the right plot, highlighted blue. On the left plot, cell type annotations are shown across diffusion component 1 and 3.

**D.** Temporal dynamics were computed using CellRank's RealTimeKernel and PsuedotimeKernel. Black and yellow dots denote starting and finishing points, respectively.

**E.** Flow model of lineage commitment of AT1 (on the left) and ciliated (on the right) cells across experimental time points, x-axis.

**F-G.** Macrostates were identified with the Generalized Perron Cluster Cluster Analysis (GPCCA) estimator (45). For each macrostate, the 50 cells most strongly associated are displayed. Macrostate distributions based on original identity (experimental time points) and cell type are shown.

**H.** Circular projections of fate probabilities towards the four terminal macrostates (AT2_1, Secretory, Basal_1, Ciliated_1). Cells are highlighted by cell type. I) Circular projection highlighting initial state forward probability of both initial macrostates (left), only AT1 (middle), and only ciliated_2 (right).

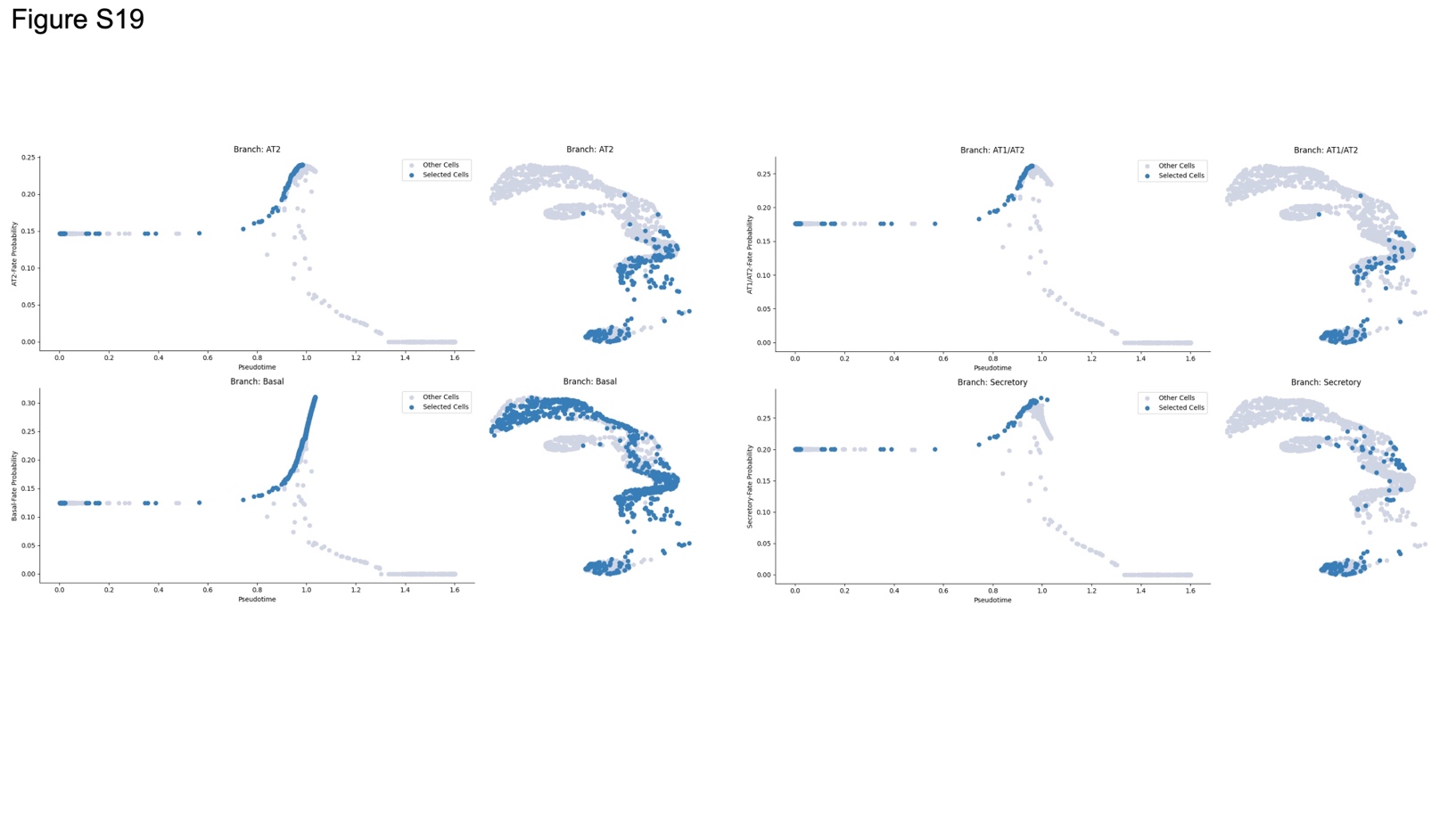

**Figure S19.** **Visualizing fate probabilities.**

(Left plot) Cell fate probabilities for the selected cell types are plotted on the y-axis and compared across psuedotime, x-axis. (Right plot) UMAP with cells highlighted that are committed to the respective lineage. Blue = member of indicated lineage, grey = not member of indicated lineage.
